# miRspongeR: an R/Bioconductor package for the identification and analysis of miRNA sponge interaction networks and modules

**DOI:** 10.1101/507749

**Authors:** Junpeng Zhang, Lin Liu, Taosheng Xu, Yong Xie, Chunwen Zhao, Jiuyong Li, Thuc Duy Le

**Affiliations:** School of Engineering, Dali University, Dali, Yunnan, 671003, China; School of Information Technology and Mathematical Sciences, University of South Australia, Mawson Lakes, SA 5095, Australia; Institute of Intelligent Machines, Hefei Institutes of Physical Science, Chinese Academy of Sciences, Hefei, 230031, China

**Author notes:** Corresponding author Email addresses: JZ LL TX YX CZ JL TDL.

**Keywords:** miRNA, ceRNA, miRNA sponge, miRNA sponge interaction networks, miRNA sponge modules, human breast invasive carcinoma

## Abstract

**Background:** A microRNA (miRNA) sponge is an RNA molecule with multiple tandem miRNA response elements that can sequester miRNAs from their target mRNAs. Despite growing appreciation of the importance of miRNA sponges, our knowledge of their complex functions remains limited. Moreover, there is still a lack of miRNA sponge research tools that help researchers to quickly compare their proposed methods with other methods, apply existing methods to new datasets, or select appropriate methods for assisting in subsequent experimental design.

**Results:** To fill the gap, we present an R/Bioconductor package, *miRspongeR*, for simplifying the procedure of identifying and analyzing miRNA sponge interaction networks and modules. It provides seven popular methods and an integrative method to identify miRNA sponge interactions. Moreover, it supports the validation of miRNA sponge interactions and the identification of miRNA sponge modules, as well as functional enrichment and survival analysis of miRNA sponge modules.

**Conclusions:** This package enables researchers to quickly evaluate their new methods, apply existing methods to new datasets, and consequently speed up miRNA sponge research.

## Background

MicroRNAs (miRNAs) are small non-coding RNAs with ~22 nucleotides. They usually induce repression or translational inhabitation of target genes through partial complementarities with multiple miRNA response elements (MREs) of the target RNA transcripts [1]. Previous studies [2, 3] have shown that miRNAs are involved in a broad range of biological processes, such as cell cycle control, cell apoptosis, cell differentiation and a diverse range of human cancers.

The competing endogenous RNA (ceRNA) hypothesis [4] considers that all types of coding and non-coding RNA transcripts may crosstalk with each other by sharing common miRNAs. By competing with each other, those ceRNAs (also known as miRNA sponges or decoys), including long non-coding RNAs (lncRNAs), pseudogenes, mRNAs and circular RNAs (circRNAs), can release parental target mRNAs from miRNAs’ control [5]. The ceRNA hypothesis challenges the traditional knowledge that coding RNAs only act as targets of miRNAs, and it provides a starting point for the investigation of the biological functions and mechanisms of miRNA sponges.

Recently, several lncRNAs, pseudogenes, circRNAs and mRNAs acting as miRNA sponges have been experimentally identified and confirmed, together with their important biological functions. For example, linc-MD1, a muscle-specific lncRNA, activates muscle-specific gene expression by regulating the expression of MAML1 and MEF2C via sponging miR-133 and miR-135 [6]. PTENP1, a pseudogene of the PTEN tumor suppression gene, can act as a sponge of PTEN-targeting miRNAs to play a growth-suppressive role [7]. As for circRNAs, CDR1as/ciRS-7 is encoded in the genome antisense to the human CDR1 (gene) locus, and affects the activity of miR-7 target genes by sponging miR-7 [8, 9]. Tay *et al*. [10] have validated two mRNAs (CNOT6L and VAPA) as miRNA sponges that regulate tumor suppressor gene PTEN, antagonise PI3K/AKT signalling, and show concordant expression patterns and copy number loss with PTEN in human cancers.

Although an increasing number of miRNA sponges have been discovered, the wet lab experiment approach for finding them is very time consuming and involves high cost. Thus computational methods [10–18] have been proposed to study miRNA sponges. For the identification of miRNA sponge interactions, the two principles, significant common miRNAs at sequence level and positively correlated at expression level, are commonly used in several computational methods. Based on the identified miRNA sponge interaction network, the miRNA sponge modules are identified by using network clustering algorithms. Here, a miRNA sponge module is defined as a cluster of miRNA sponges. It has been shown that there is a great potential to reveal the biological mechanisms in cancer by studying miRNA sponge networks and modules [5, 11, 19], and the computational methods for identifying miRNA sponge networks and modules can help broaden our study of miRNA sponges and generate hypotheses for wet lab experiments.

Recently, some tools have been developed, including spongeScan [20], miRNAsong [21], CircInteractome [22], miRNACancerMAP [23], and JAMI [24], to study miRNA sponges. Based on the sequence data of miRNAs and RNA transcripts, the first three web-based tools, spongeScan, miRNAsong and CircInteractome, regard RNA transcripts containing multiple miRNA binding sites as potential miRNA sponges. The principle of the three tools is the same with the *miRNA Homology* (*miRHomology*) [12, 13] method. The miRNACancerMAP tool uses the *positive correlation* (*pc*) method [14, 15] to identify lncRNA related miRNA sponge networks. The JAMI tool is a fast implementation of the *hermes* method [17] for inferring miRNA sponge networks.

Although these tools have been successfully applied to the study of miRNA sponges, there is still a lack of software tools for systematically identifying and analyzing miRNA sponge interaction networks and modules. Specifically, it is still difficult for researchers to choose suitable computational methods to identify miRNA sponge interactions or modules in their own data, or compare the performance of their newly developed methods with existing methods. Moreover, it is time consuming for both bioinformaticians and biologists to obtain existing algorithms for their studies as there is not a tool which contains a comprehensive collection of the algorithms. To fill this gap, we have developed an R/Bioconductor package, *miRspongeR*, to provide a pipeline for the identification and analysis of miRNA sponge interaction networks and modules. The advantage of the *miRspongeR* package is that it enables bioinformaticians and biologists to quickly compare their proposed methods with existing methods, apply existing methods to their own data, and select suitable methods for assisting in subsequent experimental design.

## Implementation

### Pipeline of miRspongeR

The *miRspongeR* package is written in R. The main pipeline of *miRspongeR* is given in Figure 1, which can be separated into three components for the identification and analysis of miRNA sponge interaction networks and modules:

- Identification of miRNA sponge interactions
- Identification of miRNA sponge modules
- Validation and analysis

**Figure 1.**
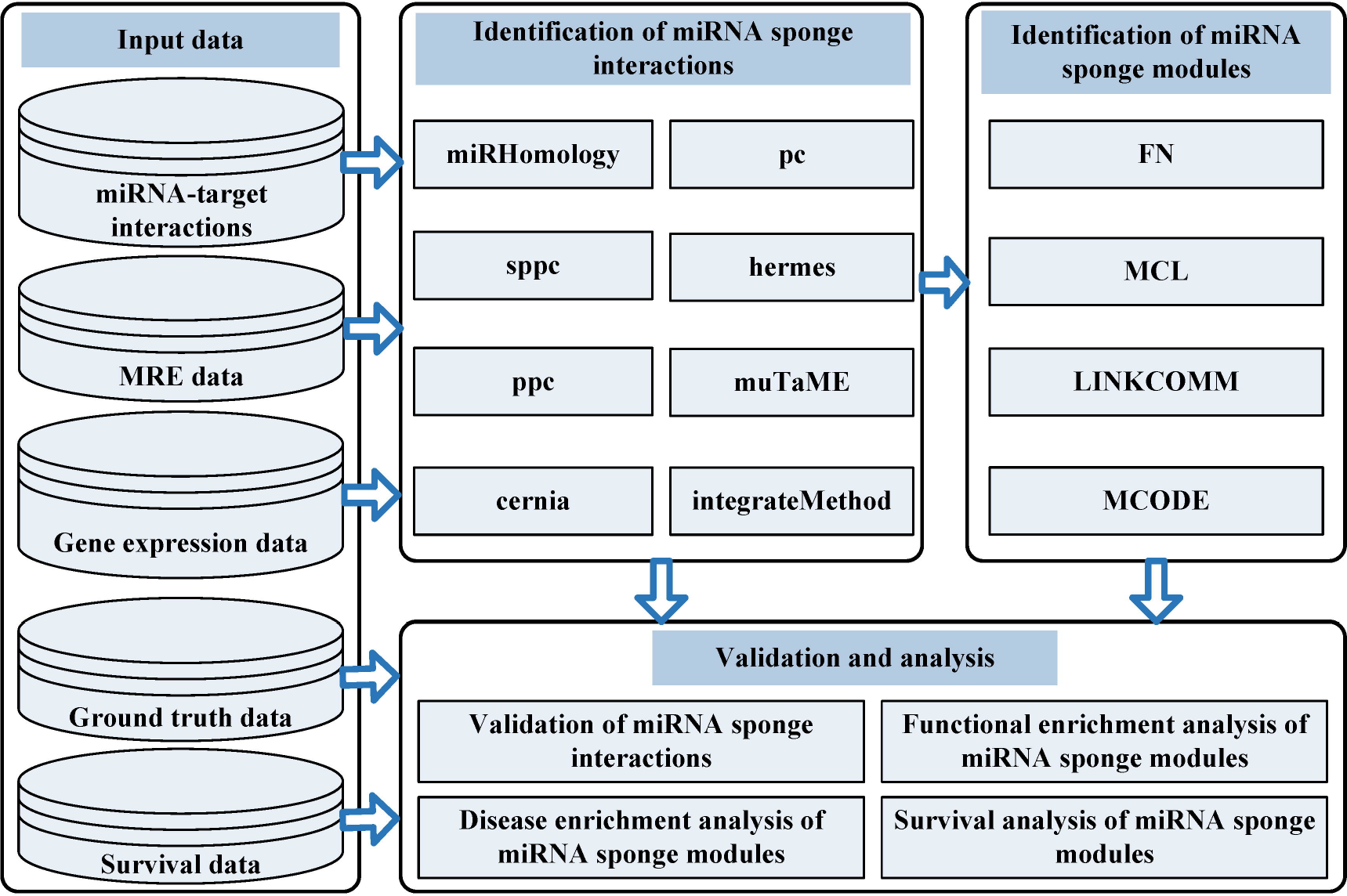
Pipeline of the miRspongeR package. The pipeline mainly contains three components: Identification of miRNA sponge interactions, Identification of miRNA sponge modules, and Validation and analysis. For the identification of miRNA sponge interactions, out of the eight implemented methods, seven (*miRHomology*, *pc*, *sppc*, *ppc*, *hermes*, *muTaME* and *cernia*) are stand-alone methods and one (*integrateMethod*) is an ensemble method which integrates the prediction results of the component methods. To understand the module-level properties of miRNA sponges, four module identification methods (FN, MCL, LINKCOMM and MCODE) are provided to identify miRNA sponge modules based on the identified miRNA sponge interaction networks. For the validation of miRNA sponge interactions, the ground truth data is used to validate predicted miRNA sponge interactions. Furthermore, enrichment analysis is performed to identify potential diseases, biological processes and pathways associated with miRNA sponge modules. Survival analysis is also performed to identify significant miRNA sponge modules which can distinguish the high and low risk tumor samples. Users can prepare their own datasets as the input to *miRspongeR*.

For input data, a miRNA-target interaction refers to the relationship of a miRNA and a validated or predicted target gene of the miRNA. The validated or predicted sites of the target gene that bind to the miRNA are known as the miRNA response elements. We say two genes share a miRNA or have a shared miRNA if they both are targets of the miRNA. When two miRNAs have a same target gene and the same miRNA response elements or binding sites on the target gene, we say the two miRNAs share miRNA target sites or they have a sharing of miRNA target sites.

In the following subsections, we describe the three components of the pipeline in detail.

### Identification of miRNA sponge interactions

To identify miRNA sponge interactions, *miRspongeR* provides the *spongeMethod* function which has implemented seven popular methods: *miRHomology* [12, 13], *positive correlation* (*pc*) [14, 15], *sppc* [16], *hermes* [17], *ppc* [11], *muTaME* [10], and *cernia* [18]. In the following, we describe each of the implemented methods in detail.

#### (1) The *miRHomology* method

The *miRNA Homology* (*miRHomology*) method identifies miRNA sponge interactions based on the homology of the shared miRNAs. Firstly, the *miRHomology* method generates all possible candidate RNA pairs that share a set of miRNAs based on putative (predicted or validated) miRNA-target interactions. Then, for each candidate RNA pair (RNA_*i*_ and RNA_*j*_), the hypergeometric test is used to evaluate the significance of the shared miRNAs by these two RNAs. The significance *p*-value is calculated as follows:

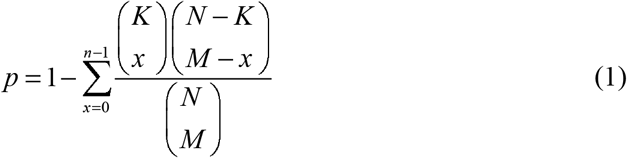

Here *N* denotes the number of all miRNAs of interest, *K* is the number of miRNAs interacting with RNA_*i*_, *M* is the number of miRNAs interacting with RNA_*j*_, and *n* is the number of the shared miRNAs by RNA_*i*_ and RNA_*j*_. The RNA-RNA pairs with significant shared miRNAs (e.g. *p*-value < 0.05) are regarded as miRNA sponge interactions.

#### (2) The *pc* method

Pearson correlation [25] is a classical method to evaluate the association between a pair of variables. The *positive correlation* (*pc*) method considers gene expression data based on the above two steps of *miRHomology* method. For the identification of miRNA sponge interactions, the correlations between each RNA pair with significant shared miRNAs are calculated. The RNA pairs with significant positive correlations (e.g. *p*-value < 0.05) are regarded as miRNA sponge interactions.

#### (3) The *sppc* method

Different from the *pc* method, the *sensitivity partial pearson correlation* (*sppc*) method considers both miRNA and mRNA expression data when inferring miRNA sponge interactions. The *sensitivity correlation* (*SC*) between each RNA pair with significant shared miRNAs is defined as follows:

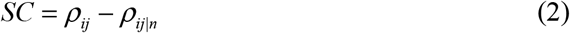

In the formula, *ρ*_ij_ is the Pearson correlation between each candidate RNA pair (RNA_*i*_ and RNA_*j*_), and *ρ*_*ij*_|_*n*_ denotes the partial Pearson correlation between the expression levels of RNA_*i*_ and RNA_*j*_, that is, the Pearson correlation between the expression levels of RNA_*i*_ and RNA_*j*_ when the expression of the *n* shared miRNAs of RNA_*i*_ and RNA_*j*_ are controlled or adjusted (to specific levels). A candidate RNA pair with high value of *SC* is more likely to be a miRNA sponge interaction.

#### (4) The *hermes* method

As Pearson correlation method only captures linear associations between variables, it cannot handle the problem of non-linear relationships well. For non-linear associations between miRNA sponges, the *hermes* method predicts miRNA sponge interactions via evidence of the sponges competing for miRNA regulation based on mutual information (MI) and conditional mutual information (CMI). For each (RNA_*i*_, miR_*k*_, RNA_*j*_) triplet, the *hermes* method calculates the statistical significance of the difference between the CMI and MI, i.e. Δ*I* = *I*[*miR_k_*; *RNA_i_*| *RNA_j_*] − *I*[*miR_k_*; *RNA_i_*] from matched miRNA and gene expression data using different CMI estimators [26]. By using Fisher’s combined probability test, the RNA pairs with statistical significance of Δ*I* (e.g. *p*-value < 0.05) are regarded as miRNA sponge interactions.

#### (5) The *ppc* method

The *partial pearson correlation* (*ppc*) method is a variant of the *hermes* method. For the linear associations between miRNA sponges, it identifies miRNA sponge interactions via evidence the sponges competing for miRNA regulation based on Pearson correlation and partial Pearson correlation. For each (RNA_*i*_, miR_*k*_, RNA_*j*_) triplet, the *ppc* method calculates the statistical significance of Δ*C* = *C*[*miR_k_*; *RNA_i_*| *RNA_j_*] − *C*[*miR_k_*; *RNA_i_*] from matched miRNA and gene expression data, where *C* stands for Pearson correlation. By using Fisher’s combined probability test, we regard the RNA pairs with statistical significance of Δ*C* (e.g. *p*-value < 0.05) as miRNA sponge interactions.

#### (6) The *muTaME* method

For a putative (validated or predicted) miRNA-target interaction, the target gene contains multiple validated or predicted MREs to bind with the miRNA. Considering the influence of MREs information in identifying miRNA sponge interactions, the *muTaME* method is implemented based on the following four scores:

- The fraction of the shared miRNAs
- The density of the MREs for all shared miRNAs
- The distribution of MREs of the putative RNA-RNA pairs
- The proportion between the overall number of MREs for a putative miRNA sponge compared with the number of miRNAs that yield these MREs

By adding the logarithm of the four scores together, we obtain the combined score for a candidate RNA pair (RNA_*i*_ and RNA_*j*_). To compare the strength of the pairs, we normalize their combined scores to obtain normalized scores in the range of [0 1], and the RNA pairs ranked high according to their scores (e.g. top 10%) are regarded as miRNA sponge interactions.

#### (7) The *cernia* method

The *cernia* method is an updated version of the *muTaME* method. It is implemented based on the following seven scores:

- The fraction of the shared miRNAs
- The density of the MREs for all shared miRNAs
- The distribution of MREs of the putative RNA-RNA pairs
- The proportion between the overall number of MREs for a putative miRNA sponge compared with the number of miRNAs that yield these MREs
- The density of the hybridization energies related to MREs for all common miRNAs
- The DT-Hybrid recommendation score
- The Pearson correlation between putative RNA-RNA pair expression data

Similarly, the method obtains a combined score by adding the logarithm of these seven scores, then normalizes the combined scores to be in the range of [0 1]. The RNA pairs with high scores (e.g. top 10%) are regarded as miRNA sponge interactions.

#### (8) The integrative method

The above seven methods each have its own merit due to different evaluating indicators. Thus, *miRspongeR* also provides the *integrateMethod* function to obtain high-confidence miRNA sponge interactions from the predictions made by different methods using majority voting. Specifically, for the predicted miRNA sponge interactions made by the above seven computational methods, we only retain those miRNA sponge interactions that are predicted by at least *k* (e.g. 3) computational methods. Certainly, users can change the setting to use more tools in the voting. To help users further understand the common and different characteristics of eight miRNA sponge interactions identification methods, we have made a summary of them in Table 1. If an input dataset contains expression levels of genes which change across different biological conditions, we say the data is dynamic; otherwise it is static.

Therefore, depending on the type of the data, i.e. dynamic or static, used by an individual method, the interactions can be divided into two types: dynamic and static. The *integrateMethod* approach contains both dynamic and static interactions, thus the type of interactions is hybrid. According to problem of linear or non-linear relationships for individual method, the type of interactions can also be divided into two types (linear and non-linear). Since the *integrateMethod* approach contains both linear and non-linear interactions, the type of interactions is also hybrid. According to the specific needs, users can select a reasonable method for their own. Moreover, regarding choosing the method to achieve the best results and performance metrics, for the same input data, the best method for the identification of miRNA sponge interactions is the one that identifies the largest percentage of experimentally validated miRNA sponge interactions.

### Identification of miRNA sponge modules

Since modularity is an important property of many biological networks, the discovery and analysis of modules in biological networks have attracted much attention within the bioinformatics community. To understand the module-level properties of miRNA sponges, we implement the *netModule* function to identify miRNA sponge modules from miRNA sponge interaction networks. Here, a miRNA sponge module denotes a cluster of miRNA sponges.

Users can select the FN [27], MCL [28], LINKCOMM [29] or MCODE [30] method for module identification. FN is a fast hierarchical agglomeration algorithm, and it is implemented by greedily optimizing the modularity for detecting community structures [27]. MCL relies on the Markov cluster algorithm, and identifies modules in biological networks by a mathematical bootstrapping procedure. The procedure simulates random walks within a biological network by the alternation of two operators called *expansion* and *inflation* [28]. LINKCOMM is a module identification method from the *linkcomm* R package [29]. It clusters the links between nodes rather than clustering nodes to identify modules. The identified modules are allowed to have same nodes, consequently uncovering the overlapping and dense community of a biological network. MCODE identifies densely connected modules based on vertex weighting (local neighbourhood density) and outward traversal (from a locally dense seed node to isolate the dense regions) [30].

### Validation and analysis

Since the ground truth of miRNA sponge interactions is still limited, it is hard to validate predicted miRNA sponge interactions generated by computational methods.

In *miRspongeR*, we provide the *spongeValidate* function to validate predicted miRNA sponge interactions by using curated miRNA sponge interactions as a source. As for the predicted miRNA sponge interactions that are not included in the source for validation, users need further wet lab experiments to know if the predictions are correct or not. The ground truth of miRNA sponge interactions is from miRSponge [31], LncCeRBase [32], LncACTdb v2.0 [33], and the three manually curated literatures [5, 11, 34]. We include the ground truth as a file in the package, and will continually update it for accurately evaluating and comparing different methods. Users can also use their own ground truth or wet lab experiments to validate predicted results.

Furthermore, to understand potential diseases, biological processes and pathways associated with miRNA sponge modules, we provide two functions, *moduleDEA* and *moduleFEA* for functional enrichment analysis. Firstly, we implement the *moduleDEA* function to make disease enrichment analysis of miRNA sponge modules. The disease databases used include Disease Ontology database (DO, http://disease-ontology.org/), DisGeNET database (DGN, http://www.disgenet.org/) and Network of Cancer Genes database (NCG, http://ncg.kcl.ac.uk/). Moreover, the *moduleFEA* function is implemented to conduct functional GO, KEGG and Reactome enrichment analysis of miRNA sponge modules. The ontology databases used are Gene Ontology database (GO, http://www.geneontology.org/), Kyoto Encyclopedia of Genes and Genomes Pathway Database (KEGG, http://www.genome.jp/kegg/), and Reactome Pathway Database (Reactome, http://reactome.org/).

Since the survival analysis can indicate whether the miRNA sponges in the discovered miRNA sponge modules are good predictors of the metastasis risks of cancer patients or not, it can provide us the clue that the miRNA sponges may be associated with and potentially contributing to the metastasis or survival of cancer patients. In *miRspongeR*, we implement the *moduleSurvival* function to perform the survival analysis. A multivariate Cox model is used to calculate the risk score of a sample. All the samples are divided into the high risk and the low risk groups according to their risk scores, and the Log-rank test is used to test for the difference between groups. The Hazard Ratio (HR) between the high and the low risk groups is also calculated.

## Application

In this section, we apply the *miRspongeR* package to the BRCA dataset. The matched Level 3 IlluminaHiSeq miRNA and mRNA expression data of human breast invasive carcinoma (BRCA) is obtained from Paci *et al*. [16]. The gene-level expression values are in Fragments Per Kilobase Million (FPKM) units. Paci *et al*. obtained the expression data from The Cancer Genome Atlas (TCGA, https://cancergenome.nih.gov/), and restricted the data to 72 individuals where the tumor and normal tissues were from the same BRCA patients. A miRNA or mRNA with missing values in more than 10% of the samples is removed. The remaining missing values are imputed using the *k*-nearest neighbours (KNN) algorithm in the *impute* R package [35]. We use the *limma* R package [36] to infer differentially expressed miRNAs and mRNAs between tumour and normal samples. After the analysis, we identify 161 miRNAs and 5370 mRNAs which are differentially expressed at a significant level (adjusted *p*-value < 1E-04, adjusted by Benjamini & Hochberg method). The survival data of the 72 BRCA tumour samples is obtained from TCGA.

To obtain high-quality candidate miRNA sponge interaction pairs, we use experimentally validated miRNA-target interactions from miRTarBase v7.0 [37] for the case study. We are only interested in the miRNA-target interactions supported by strong experimental evidences (Reporter assay or Western blot) in miRTarBase. Between 161 miRNAs and 5370 mRNAs which are differentially expressed, we obtain 1251 unique miRNA-target interactions from miRTarBase. The MREs information is obtained from Sardina *et al*. [18]. They employ miRanda algorithm [38] to predict binding sites, and MREs have the hybridization energy values no more than 0 are reserved. In total, we have a list of 824828 MREs related to the differentially expressed miRNAs and mRNAs. By using the default ground truth in the *miRspongeR* package, we have a list of 1465 unique validated miRNA sponge interactions in total. For the identification of miRNA sponge interactions, the cutoffs to be set by users are recommended as follows: 0.05 for the significant *p*-value cutoff of the shared miRNAs, 0.05 for the *p*-value cutoff of significant positive correlations for the *pc* method, and 0.1 for the *sensitivity correlation* cutoff for the *sppc* method. A higher value of the *sensitivity correlation* cutoff will generate a smaller number of miRNA sponge interactions. As for the *hermes* and *ppc* methods, we suggest use 0.05 as the statistical significance *p*-value cutoff of evaluating Δ*I* and Δ*C*. All the *p*-values above are adjusted by the Benjamini & Hochberg method, and a smaller *p*-value cutoff will identify a smaller number of miRNA sponge interactions. The number of permutations for both the *hermes* and *ppc* methods has been set to 100 by default in the package. The cutoff of normalized score is set to 0.5 by default in the package for the *muTaME* and *cernia* methods. For the *integrateMethod* method, we only retain the predicted miRNA sponge interactions which are predicted by at least 3 out of the 7 component methods.

For the identification miRNA sponge modules in the BRCA dataset, the module size cutoff for the FN, MCL, LINKCOMM and MCODE methods is set to 3 by default in the package. Moreover, we conduct enrichment and survival analysis of the identified miRNA sponge modules by using default settings in the package.

## Results

The *miRspongeR* package is developed for the identification and analysis of miRNA sponge interaction networks and modules. In this section, we show the usages of the *miRspongeR* package by conducting a case study. For the case study, the BRCA dataset used can be obtained from https://github.com/zhangjunpeng411/miRspongeR_dataset and the running scripts can be seen in Additional file 1.

In the case study, we only focus on studying mRNA related miRNA sponge interaction networks and modules. It is noted that *miRspongeR* can be applied to study the miRNA sponge interaction networks and modules involving other types of RNA (lncRNA, pseudogene, circRNA, etc.). For instance, if we want to consider lncRNAs acting as miRNA sponges, the input miRNA-target interactions and MRE data to *miRspongeR* should contain miRNA-lncRNA interactions and the gene expression data should include matched lncRNA expression data.

### Identification and validation of BRCA-related miRNA sponge interactions

In this section, we use the *miRspongeR* package, including the 7 methods introduced in the section *Identification of miRNA sponge interactions* and the integrated method (*integrateMethod*) to identify BRCA-related miRNA sponge interactions.

As shown in Figure 2, the numbers of predicted miRNA sponge interactions by the 7 individual methods are different. The number of predicted miRNA sponge interactions shared by the 7 methods is 5. Specifically, the *miRHomology* method independently predicted 44 miRNA sponge interactions which are not predicted by any of the other methods. The reason is that the *miRHomology* method only uses a commonly used constraint (significant sharing of common miRNAs) to identify miRNA sponge interactions. By using the *integrateMethod* method, we obtain a list of 87 high-confidence miRNA sponge interactions. The detailed information of predicted miRNA sponge interactions by the methods can be seen in Additional file 2.

**Figure 2.**
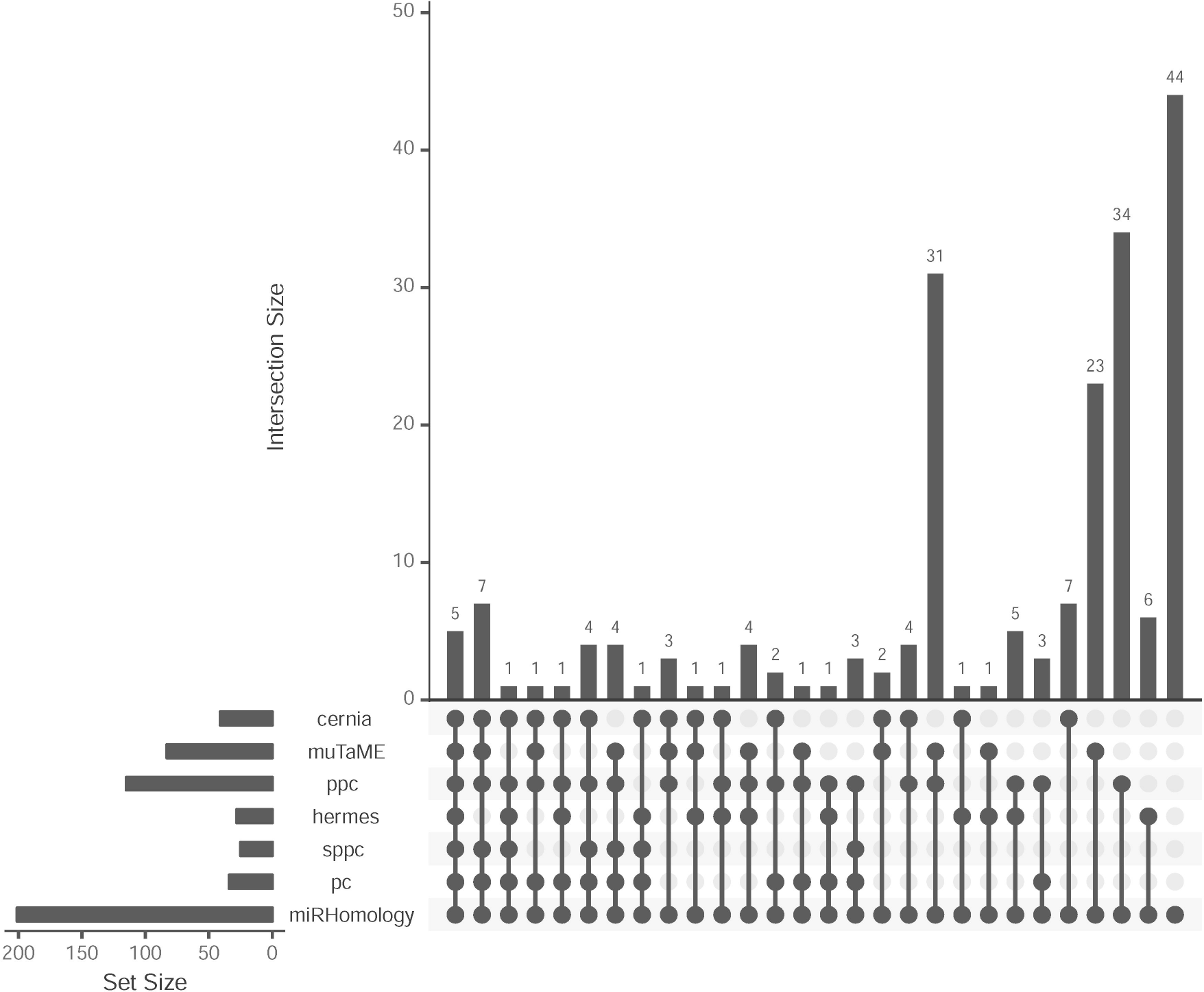
UpSet plot [39] to show the intersections between predicted miRNA sponge interactions by the 7 built-in individual methods. Each column corresponds to an exclusive intersection that includes the elements of the sets denoted by the dark circles, but not of the others. The intersection size between different methods represents exclusive intersections, i.e. the intersection set not in a subset of any other intersection set.

To investigate the overlap of any 2 of 7 individual methods, we further use the Venn diagrams to show pair-wise comparison of overlapping results for the 7 individual methods. The detailed information of pair-wise comparison results can be seen in Additional file 3.

Furthermore, we use the ground truth to validate the predicted miRNA sponge interactions. As a result, for all the methods only 1 miRNA sponge interaction, (PTEN:ZEB1, where “:” denotes competing relationship), is validated by the ground truth. In terms of the number of experimentally validated miRNA sponge interactions, these methods perform the same. However, in terms of the percentage of experimentally validated miRNA sponge interactions, the *sppc* method performs the best.

### Module identification from BRCA-related miRNA sponge interaction network

Since the *integrateMethod* method integrates the results of 7 individual methods (*miRHomology*, *pc*, *sppc*, *hermes*, *ppc*, *muTaME* and *cernia*) to infer high-confidence miRNA sponge interaction network, in this section, we focus on module identification from BRCA-related miRNA sponge interaction network discovered by this integrated method. As shown in Table 2, the four module identification methods (FN, MCL, LINKCOMM and MCODE) obtain different results of the identified miRNA sponge modules. As LINKCOMM allows overlapping nodes between modules, it identified the largest number of miRNA sponge modules.

### Enrichment and survival analysis of BRCA-related miRNA sponge modules

In this section, we conduct enrichment and survival analysis of the identified BRCA-related miRNA sponge modules described in the previous section. Here, if a term (DO, DGN, NCG, GO, KEGG or Reactome term) is enriched in a miRNA sponge module with adjusted *p*-value < 0.05 (adjusted by Benjamini & Hochberg method), the term is a significantly enriched term. If a miRNA sponge module can distinguish the high and the low risk tumor samples (Log-rank *p*-value < 0.05, HR > 1.5), the miRNA sponge module is regarded as a significant miRNA sponge module.

The overall number of significantly enriched terms can be seen in Figure 3. Overall, the identified modules by the LINKCOMM method have the largest number of significantly enriched DGN, GO and Reactome terms. The identified modules by MCODE have the largest number of significantly enriched DO and NCG terms. The modules identified by the FN method have the largest number of significantly enriched KEGG terms. For each method, the number of significantly enriched terms in each module can be seen in Additional file 4. Due to a lack of comprehensive knowledge of enriched terms indirectly and directly associated with BRCA, we use all significantly enriched terms (DO, DGN, NCG, GO, KEGG and Reactome terms) associated with the miRNA sponge modules. It is noted that users can further investigate these significantly enriched terms in the context of BRCA.

**Figure 3.**
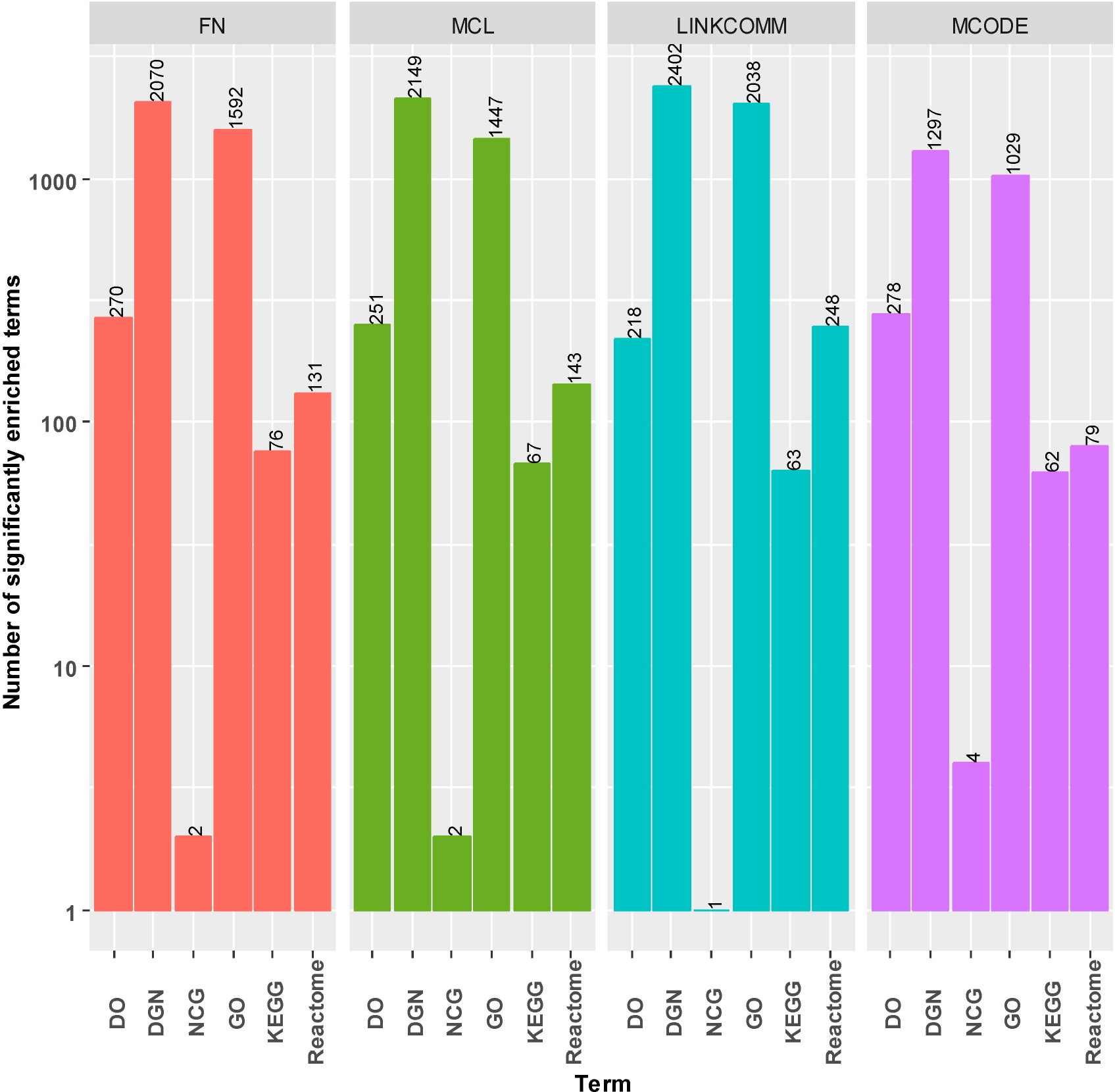
Overall number of significantly enriched terms in BRCA-related miRNA sponge modules using FN, MCL, LINKCOMM and MCODE. The significantly enriched terms include DO, DGN, NCG, GO, KEGG and Reactome terms.

As shown in Table 3, the numbers of significant miRNA sponge modules for distinguishing the high and the low risk BRCA samples are 4, 2, 5 and 2 for FN, MCL, LINKCOMM and MCODE methods, respectively. This result indicates that LINKCOMM method identifies the largest number of significant miRNA sponge modules.

## Discussion

*miRspongeR* has been released under the GPL-3.0 License, and is available at http://bioconductor.org/packages/miRspongeR/ and https://github.com/zhangjunpeng411/miRspongeR. The user manual of *miRspongeR* provides examples illustrating the use of the functions of *miRspongeR*, and can be seen in https://bioconductor.org/packages/release/bioc/vignettes/miRspongeR/inst/doc/miRspongeR.html.

The putative miRNA-target interactions are crucial for the identification of miRNA sponge interaction. The miRNA-target identification methods, such as miRanda [38], TargetScan [40] and DIANA-microT [41], predict many false positives and the number of miRNA target genes is largely overestimated [42], affecting the accuracy of the candidate miRNA sponge interaction pairs. Therefore, instead of predicted miRNA-target interactions, we use experimentally validated miRNA-target interactions in the case study. The experimentally validated miRNA-target interactions can be used to provide high-quality candidate miRNA sponge interaction pairs. According to the practical situation, users can also use their own predicted miRNA-target interactions to obtain candidate miRNA sponge interaction pairs. Moreover, the number of candidate miRNA sponges is closely associated with the putative miRNA-target interactions used. In the case study, we only use the miRNA-target interactions supported by strong experimental evidences (Reporter assay or Western blot) in miRTarBase. The maximum number of miRNA sponges expected to be identified depends on the number of targets in the putative miRNA-target interactions. However, experimentally validated miRNA-target interactions are far from complete. A possible solution is to integrate multiple experimentally validated datasets, such as miRTarBase [37], TarBase [43] and miRWalk [44], to obtain a more comprehensive list of putative miRNA-target interactions.

## Conclusions

Recently, miRNA sponges have been demonstrated to play important roles in human cancers by modulating the expression of key oncogenes and tumor-suppressor genes. Since biological experiments generate a large amount of data associated with miRNA sponges, this creates a strong demand for data analysis in statistics computing environments such as R. In this paper, we introduce an R/Bioconductor package that enables R users to use popular computational methods for the identification and analysis of miRNA sponge interaction networks and modules. As the research into miRNA sponges is just emerging, we will continuously include new methods in the package in the future. We believe that *miRspongeR* will be a useful tool for the research of miRNA sponges.

## Availability and requirements

**Project name:** miRspongeR

**Project home page:** http://bioconductor.org/packages/miRspongeR/

**Operating system(s):** Platform independent

**Programming language:** R

**Other requirements:** R (>= 3.5.0)

**License:** GPL-3

**Any restrictions to use by non-academics:** licence needed

## Abbreviations

miRNA: microRNA
MRE: miRNA response element
ceRNA: Competing endogenous RNA
lncRNA: Long non-coding RNA
circRNA: Circular RNA
miRHomology: miRNA Homology
pc: positive correlation
sppc: sensitivity partial pearson correlation
SC: Sensitivity correlation
MI: Mutual information
CMI: Conditional mutual information
ppc: partial pearson correlation
DO: Disease ontology
NCG: Network of Cancer Genes
GO: Gene Ontology
KEGG: Kyoto Encyclopedia of Genes and Genomes Pathway
HR: Hazard Ratio
BRCA: Breast invasive carcinoma
FPKM: Fragments Per Kilobase Million
TCGA: The Cancer Genome Atlas
KNN: *k*-nearest neighbours

## Declarations

## Acknowledgements

We are grateful to the Bioconductor Project, for the valuable comments on the codes to greatly improve the *miRspongeR* package.

## Availability of data and materials

The datasets in the current study are available at https://github.com/zhangjunpeng411/miRspongeR_dataset.

## Authors’ contributions

JZ and TDL conceived the idea of this work. LL, TX and JL refined the idea. JZ designed and performed the experiments. TX, YX and CZ participated in the design of the study and performed the statistical analysis. JZ, TDL, LL, TX and JL drafted the manuscript. All authors revised the manuscript. All authors read and approved the final manuscript.

## Consent for publication

Not applicable.

## Ethics approval and consent to participate

Not applicable.

## Tables

**Table 1 – Summary of the eight methods for identifying miRNA sponge interactions.**

**Table 2 – BRCA-related miRNA sponge modules using FN, MCL, LINKCOMM and MCODE methods.**

**Table 3 – Survival analysis of BRCA-related miRNA sponge modules using FN, MCL, LINKCOMM and MCODE.**

HRlow95 and HRup95 denote the lower and upper of 95% confidence interval of HR, respectively. The identified significant miRNA sponge modules can distinguish the high and the low risk BRCA samples.

## Additional files

**Additional file 1 – The used running R scripts for reproducing the results of the case study.**

**Additional file 2 – The predicted miRNA sponge interactions by the 8 built-in methods.**

**Additional file 3 – Pair-wise comparison results of the 7 built-in individual method.**

**Additional file 4 – The number of significantly enriched terms in each BRCA-related miRNA sponge module using FN, MCL, LINKCOMM and MCODE.** The significantly enriched terms contain DO, DGN, NCG, GO, KEGG and Reactome terms.

