## Supplementary material for "miRspongeR: an R/Bioconductor package for the identification and analysis of miRNA sponge interaction networks and modules": Table 1

**Table 1 Summary of the eight methods for identifying miRNA sponge interactions**

| **Methods** | **Input** | **Type of interactions** | **Advantages/disadvantages** |
| --- | --- | --- | --- |
| miRHomology | miRNA-target interactions | static | - the number of miRNA sponges is largely overestimated - ignore gene expression data and MREs information - simple and fast |
| pc | miRNA-target interactions, gene expression data | dynamic, linear | - ignore non-linear interactions - ignore miRNA expression data and MREs information - simple and fast |
| sppc | miRNA-target interactions, gene expression data | dynamic, linear | - ignore non-linear interactions - ignore MREs information - employ sensitivity correlation to evaluate the influence of miRNAs |
| hermes | miRNA-target interactions, gene expression data | dynamic, non-linear | - ignore MREs information - time consuming - capture non-linear interactions by calculating the statistical significance of 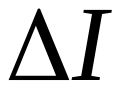 |
| ppc | miRNA-target interactions, gene expression data | dynamic, linear | - ignore non-linear interactions - ignore MREs information - time consuming - capture linear interactions by calculating the statistical significance of 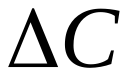 |
| muTaME | miRNA-target interactions, MREs | static | - ignore gene expression data - consider MREs information |
| cernia | miRNA-target interactions, gene expression data, MREs | dynamic, linear | - ignore non-linear interactions - ignore miRNA expression data - consider MREs information |
| integrateMethod | miRNA sponge interaction networks | hybrid | - contain dynamic and static interactions - include linear and non-linear interactions - high-confidence but time consuming |
