## Supplementary material for "miRspongeR: an R/Bioconductor package for the identification and analysis of miRNA sponge interaction networks and modules": Table 2

**Table 2 BRCA-related miRNA sponge modules using FN, MCL, LINKCOMM and MCODE methods**

| **Module methods** | **Module ID** | **#miRNA sponges** | **miRNA sponges** |
| --- | --- | --- | --- |
| FN | 1 | 14 | *MYB, EIF4E, E2F3, CDC25A, DNMT3A, VEGFA, BCL2, CCNE1, ETS1, DNMT3B, BIRC5, KDR, FGF2, CCND3* |
|  | 2 | 12 | *NOTCH1, ZEB1, ZEB2, ZFPM2, TCF7L1, ELMO2, KLHL20, KLF11, CCNE2, DLC1, WASF3, WDR37* |
|  | 3 | 7 | *NFIA, FOXO1, BDNF, RECK, CREB1, SNAI2, RUNX2* |
|  | 4 | 5 | *PTEN, KLF4, TIMP2, PPARA, BCL2L11* |
|  | 5 | 4 | *CDK6, MITF, KRAS, CCND2* |
|  | 6 | 3 | *SMAD4, TP53INP1, CASP3* |
| MCL | 1 | 5 | *PTEN, KLF4, TIMP2, PPARA, BCL2L11* |
|  | 2 | 12 | *NOTCH1, ZEB1, ZEB2, ZFPM2, TCF7L1, ELMO2, KLHL20, KLF11, CCNE2, DLC1, WASF3, WDR37* |
|  | 3 | 10 | *EIF4E, E2F3, CDC25A, VEGFA, BCL2, CCNE1, ETS1, BIRC5, FGF2, CCND3* |
|  | 4 | 4 | *CDK6, MITF, KRAS, CCND2* |
|  | 5 | 3 | *SMAD4, TP53INP1, CASP3* |
|  | 6 | 5 | *NFIA, FOXO1, BDNF, RECK, SNAI2* |
| LINKCOMM | 1 | 4 | *MYB, ZEB2, ZEB1, DLC1* |
|  | 2 | 5 | *ZEB1, ZEB2, ZFPM2, KLF11, WDR37* |
|  | 3 | 3 | *FOXO1, RECK, SNAI2* |
|  | 4 | 4 | *E2F3, CDC25A, CCNE1, CCND3* |
|  | 5 | 7 | *PTEN, ZEB1, ZFPM2, TCF7L1, KLHL20, KLF11, WDR37* |
|  | 6 | 3 | *CDC25A, BCL2, CCNE1* |
|  | 7 | 5 | *PTEN, TIMP2, KLF4, PPARA, BCL2L11* |
|  | 8 | 4 | *EIF4E, E2F3, BIRC5, FGF2* |
|  | 9 | 6 | *E2F3, MYB, ZEB2, CCNE2, TCF7L1, KLF11* |
|  | 10 | 4 | *NOTCH1, ZEB1, ZEB2, ETS1* |
|  | 11 | 3 | *VEGFA, E2F3, ETS1* |
|  | 12 | 7 | *ZEB1, ZEB2, ZFPM2, TCF7L1, KLHL20, KLF11, DLC1* |
| MCODE | 1 | 25 | *CDK6, NFIA, EIF4E, DNMT3A, MITF, BDNF, PIM1, KLF4, CREB1, SMAD4, SMAD3, KRAS, CCND2, TIMP2, DNMT3B, TWIST1, BIRC5, KDR, FGF2, PPARA, RUNX2, TP53INP1, CASP3, SMAD2, BCL2L11* |
|  | 2 | 24 | *MYB, PTEN, FOXO1, NOTCH1, ZEB1, ZEB2, ZFPM2, E2F3, CDC25A, TCF7L1, RECK, VEGFA, ELMO2, BCL2, KLHL20, KLF11, CCNE2, CCNE1, ETS1, DLC1, SNAI2, WASF3, WDR37, CCND3* |
