## Supplementary material for "miRspongeR: an R/Bioconductor package for the identification and analysis of miRNA sponge interaction networks and modules": Table 3

**Table 3 Survival analysis of BRCA-related miRNA sponge modules using FN, MCL, LINKCOMM and MCODE**

| **Module methods** | **Module ID** | **Chi-square** | ***p*-value** | **HR** | **HRlow95** | **HRup95** |
| --- | --- | --- | --- | --- | --- | --- |
| FN | 1 | 9.96 | 1.60E-03 | 2.75 | 1.41 | 5.38 |
|  | 2 | 8.32 | 3.92E-03 | 2.57 | 1.33 | 4.96 |
|  | 3 | 4.34 | 3.71E-02 | 1.95 | 1.00 | 3.80 |
|  | 5 | 4.72 | 2.98E-02 | 2.00 | 1.04 | 3.87 |
| MCL | 2 | 8.32 | 3.92E-03 | 2.57 | 1.33 | 4.96 |
|  | 4 | 4.72 | 2.98E-02 | 2.00 | 1.04 | 3.87 |
| LINKCOMM | 2 | 4.78 | 2.88E-02 | 2.03 | 1.04 | 3.94 |
|  | 5 | 14.67 | 1.28E-04 | 3.24 | 1.64 | 6.43 |
|  | 9 | 3.97 | 4.62E-02 | 1.94 | 1.01 | 3.73 |
|  | 10 | 5.41 | 2.00E-02 | 2.17 | 1.12 | 4.18 |
|  | 12 | 9.88 | 1.67E-03 | 2.81 | 1.45 | 5.46 |
| MCODE | 1 | 23.83 | 1.05E-06 | 4.52 | 2.26 | 9.03 |
|  | 2 | 23.84 | 1.05E-06 | 4.94 | 2.52 | 9.69 |

HRlow95 and HRup95 denote the lower and upper of 95% confidence interval of HR, respectively. The identified significant miRNA sponge modules can distinguish the high and the low risk BRCA samples.
