## Additional file 3 for "miRspongeR: an R/Bioconductor package for the identification and analysis of miRNA sponge interaction networks and modules"

We use the Venn diagrams to show pair-wise comparison of overlapping results for 7 individual methods (miRHomology, pc, sppc, hermes, ppc, muTaME and cernia). In total, we have 21 cases of pair-wise comparison in the following.

**A**

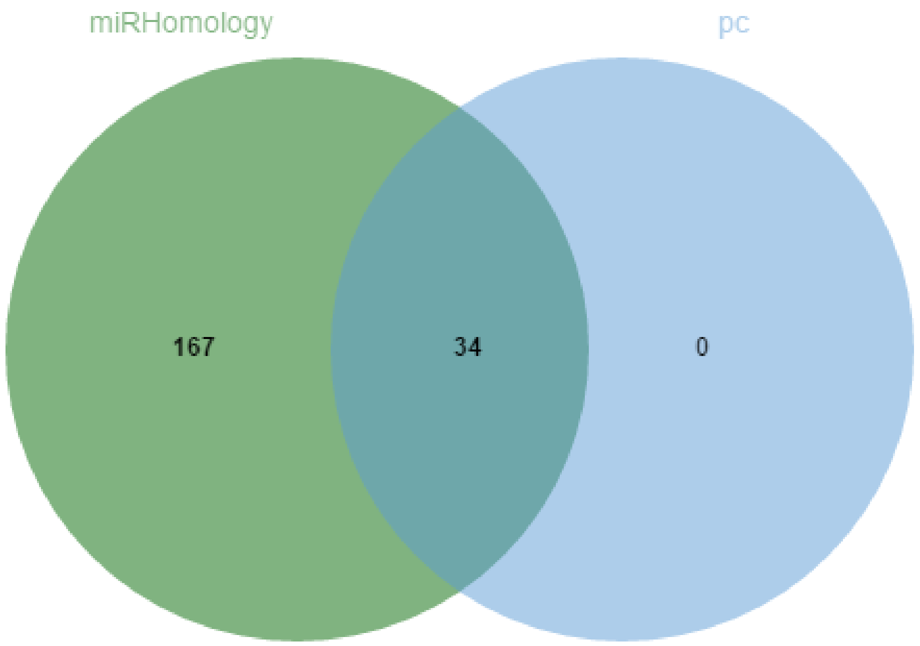


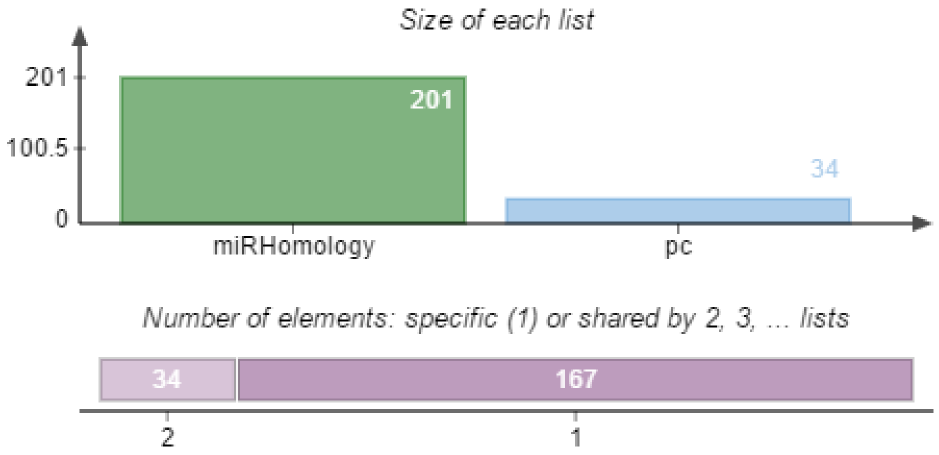

B

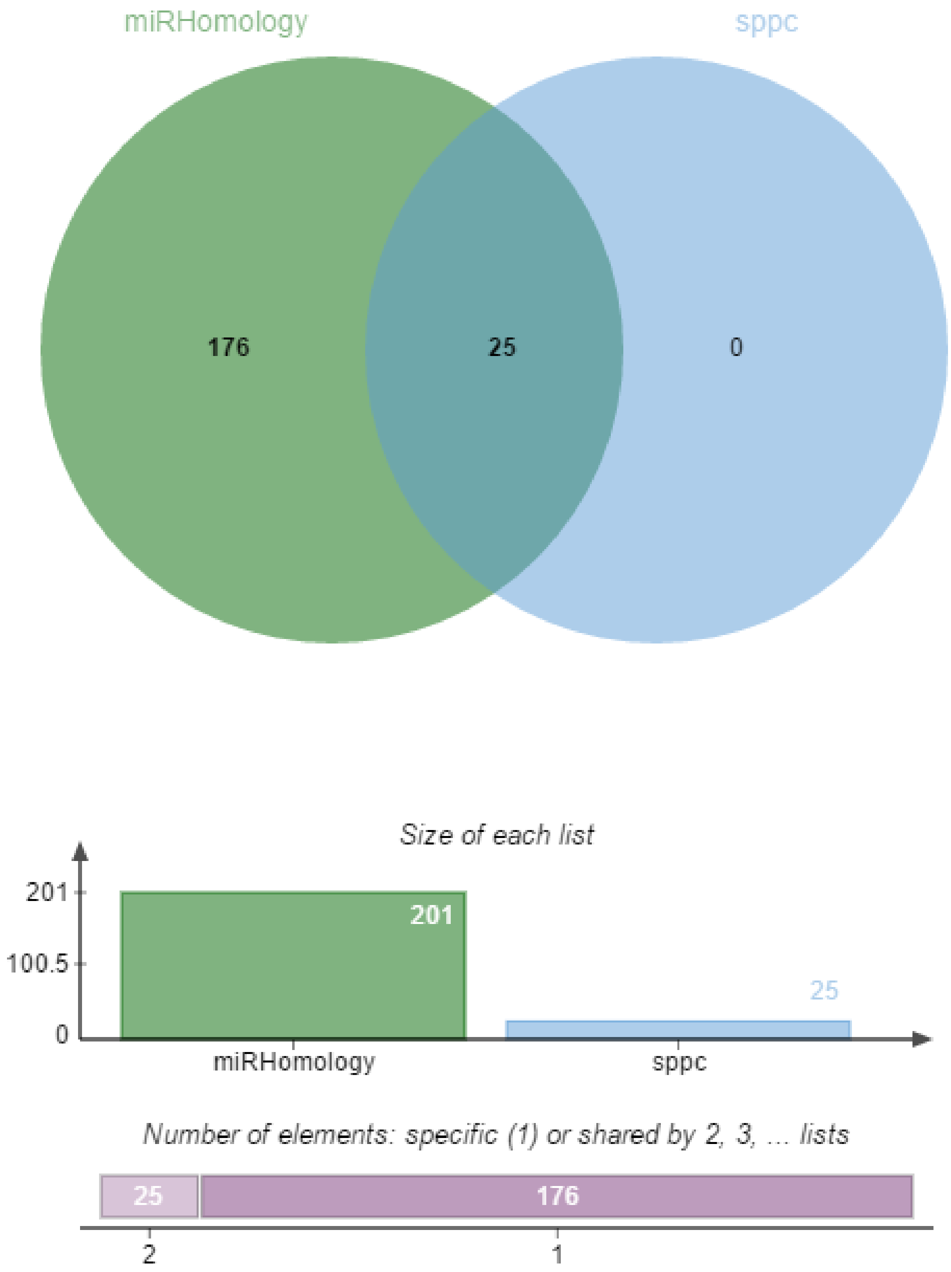
miRHomology
sppc
176
25
0
Size of each list
201
100.5
0
201
25
miRHomology
sppc
Number of elements: specific (1) or shared by 2, 3, ... lists
25
176
2
1

C

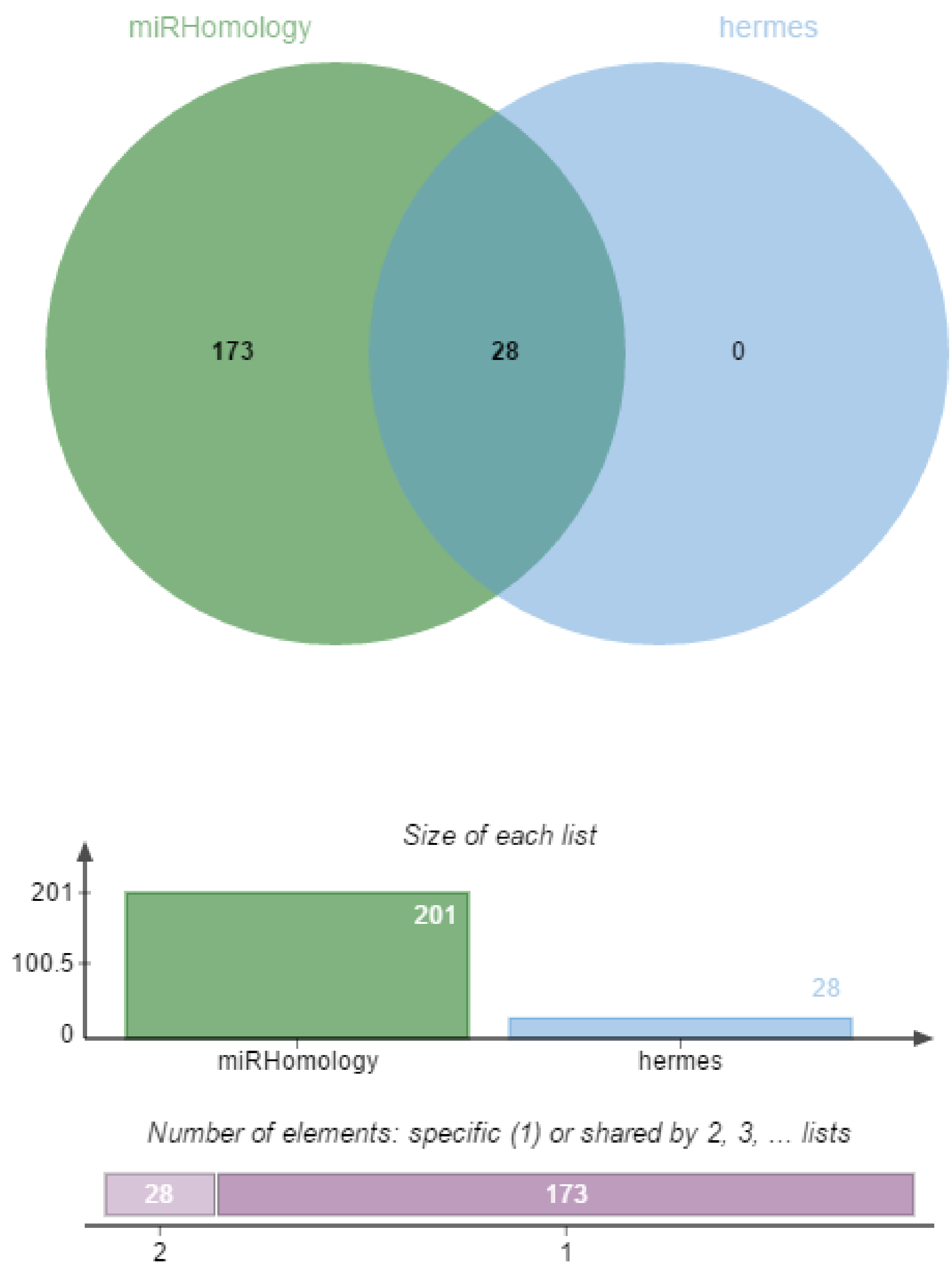
miRHomology
hermes
173
28
0
Size of each list
201
100.5
0
201
28
miRHomology
hermes
Number of elements: specific (1) or shared by 2, 3, ... lists
28
173
2
1

D

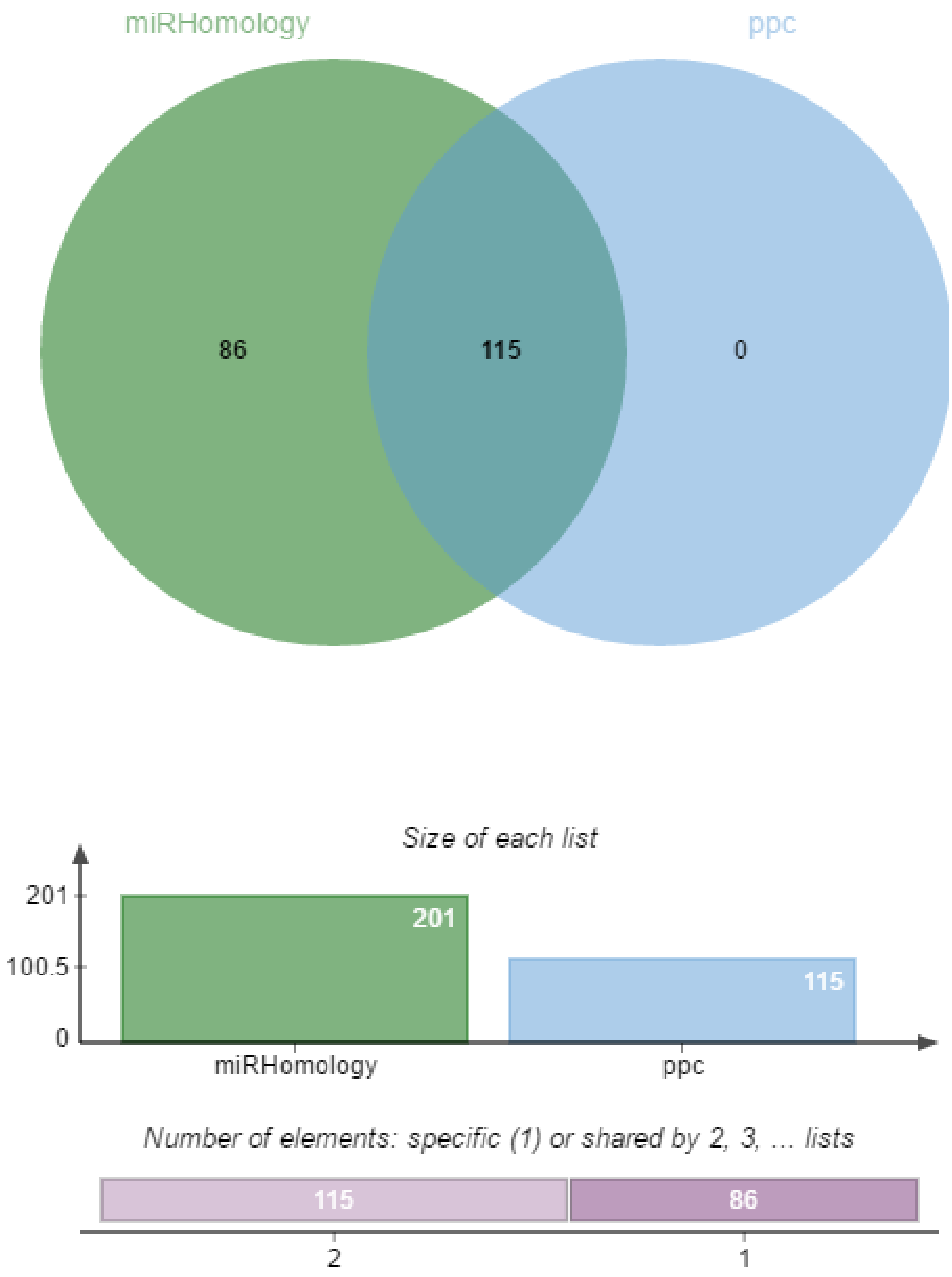

miRHomology
ppc
86
115
0
Size of each list
201
100.5
0
201
115
miRHomology
ppc
Number of elements: specific (1) or shared by 2, 3, ... lists
115
86
2
1

E

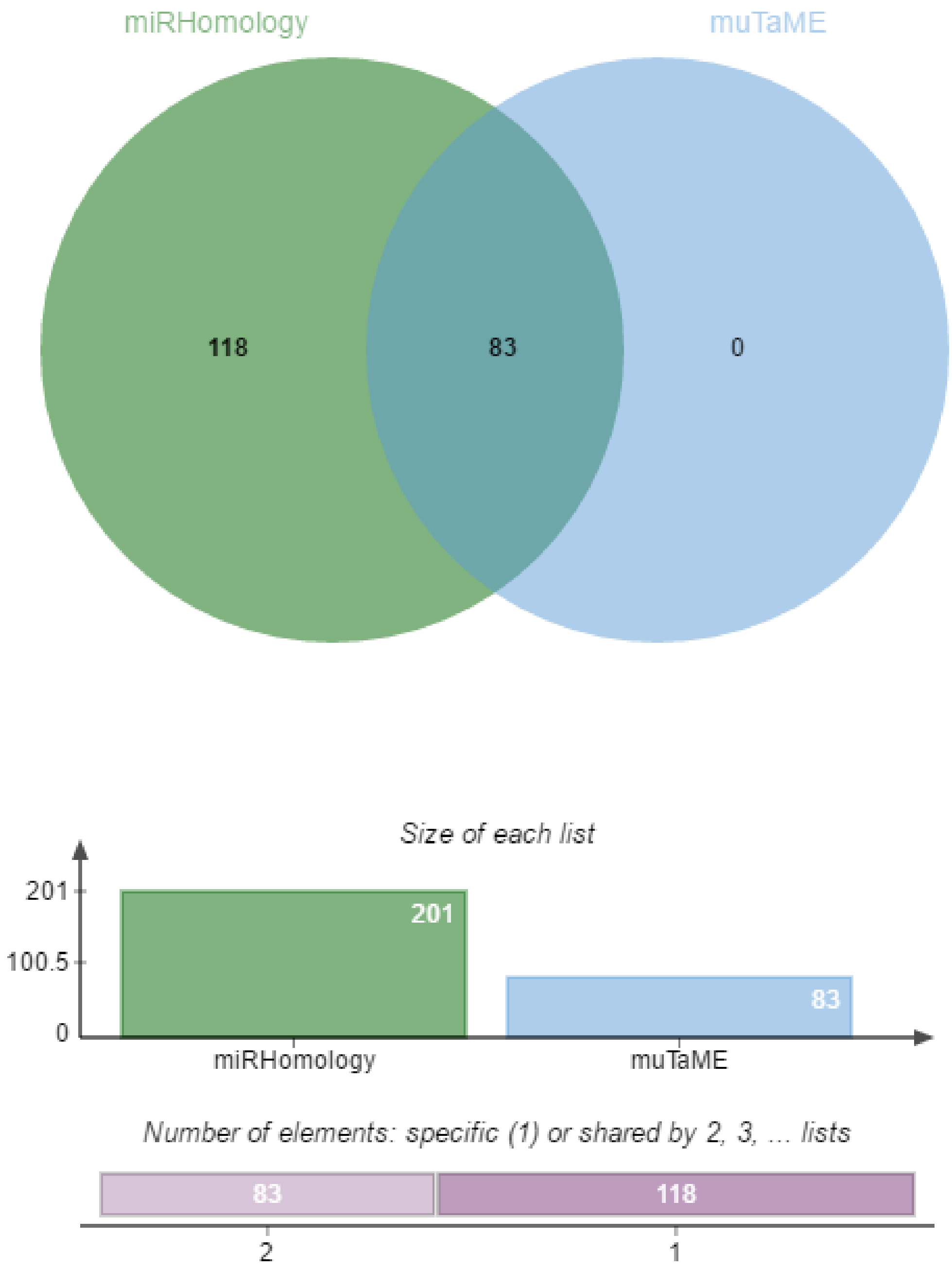

miRHomology
muTaME
118
83
0
Size of each list
201
100.5
0
201
83
miRHomology
muTaME
Number of elements: specific (1) or shared by 2, 3, ... lists
83
118
2
1

F

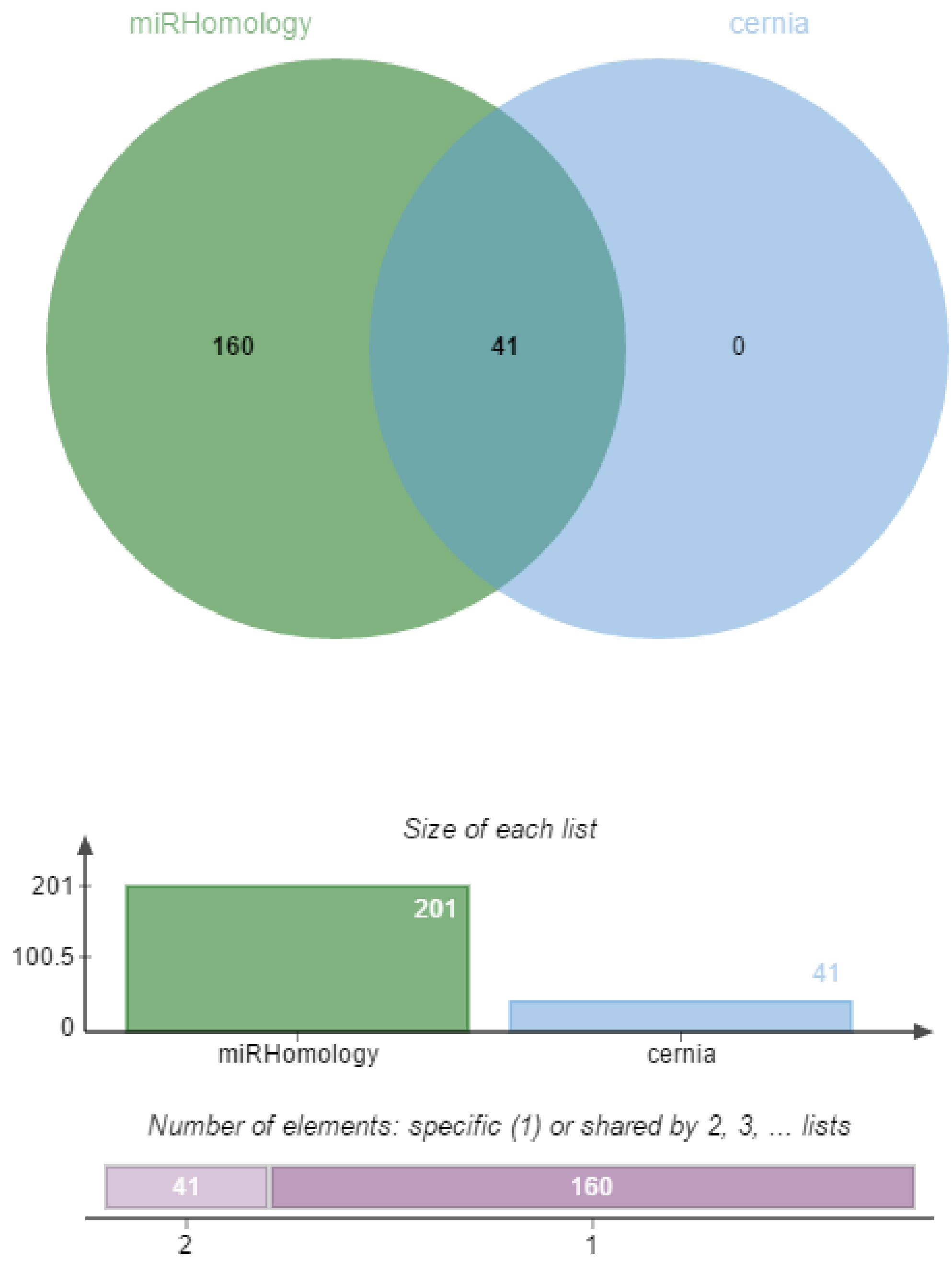

miRHomology
cernia
160
41
0
Size of each list
201
100.5
0
201
41
miRHomology
cernia
Number of elements: specific (1) or shared by 2, 3, ... lists
41
160
2
1

G

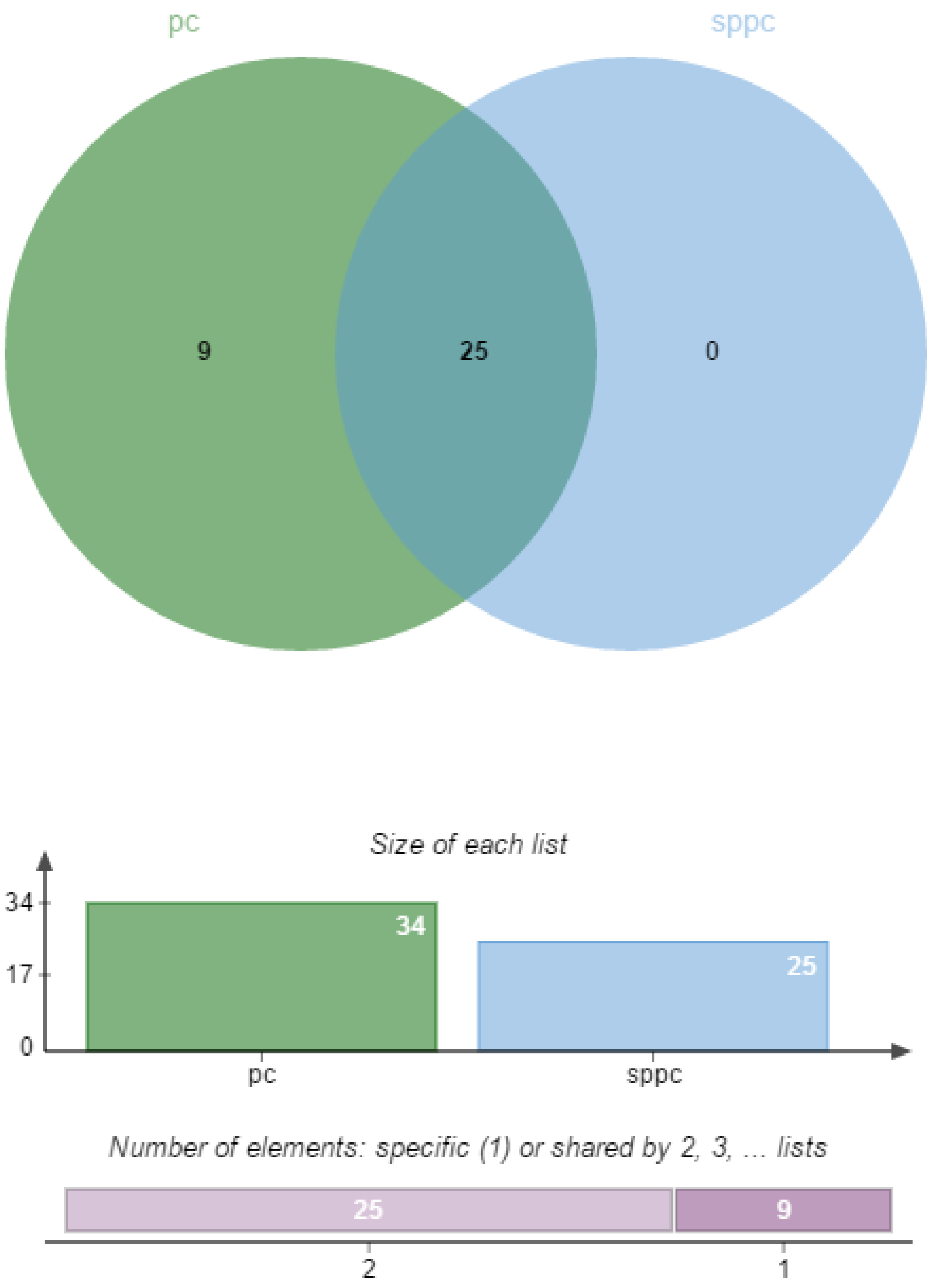
pc
sppc
9
25
0
Size of each list
34
17
0
34
25
pc
sppc
Number of elements: specific (1) or shared by 2, 3, ... lists
25
9
2
1

H

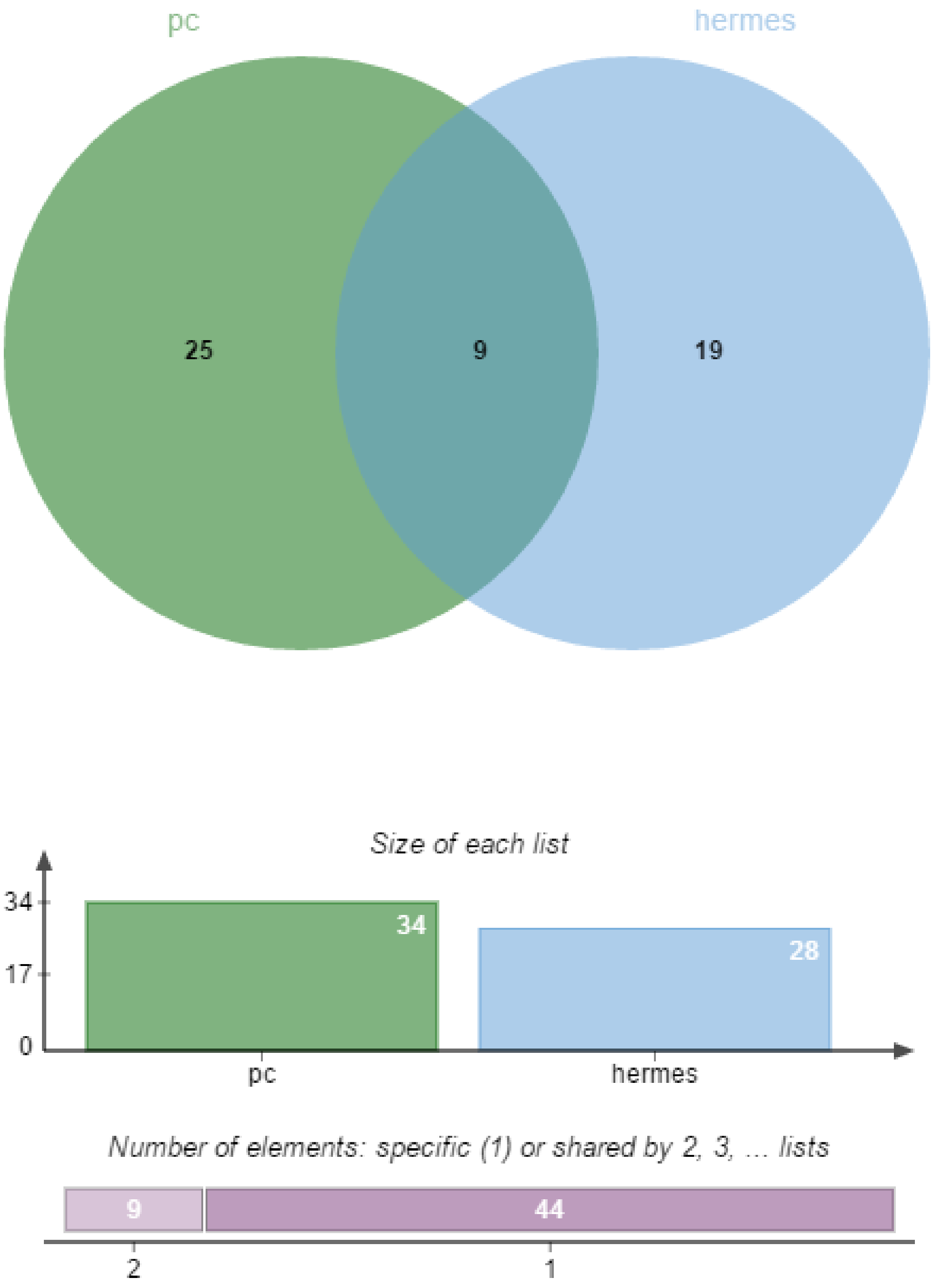

pc
hermes
25
9
19
Size of each list
34
17
0
34
28
pc
hermes
Number of elements: specific (1) or shared by 2, 3, ... lists
9
44
2
1

I

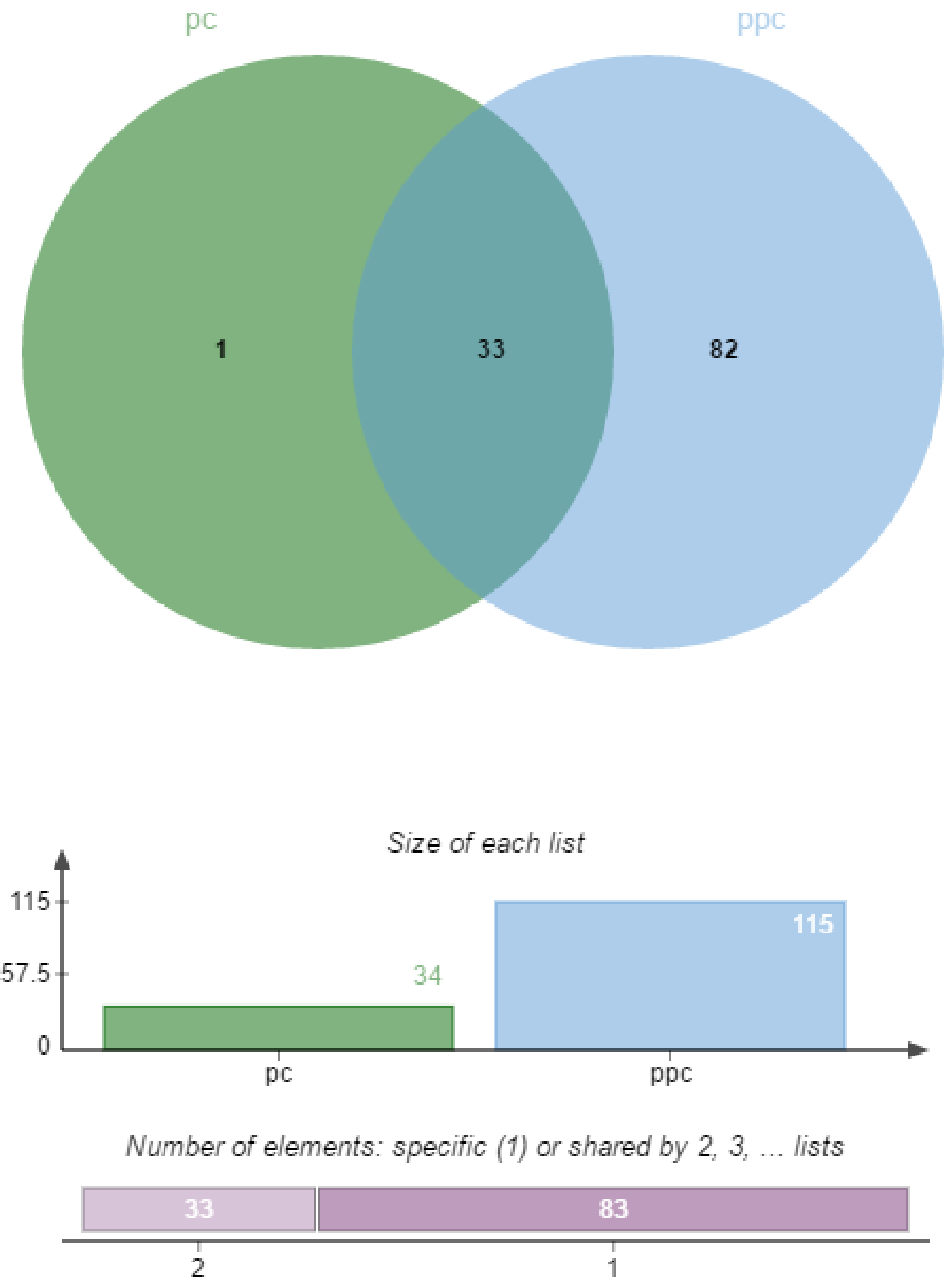
pc
ppc
1
33
82
Size of each list
115
57.5
0
34
115
pc
ppc
Number of elements: specific (1) or shared by 2, 3, ... lists
33
83
2
1

J

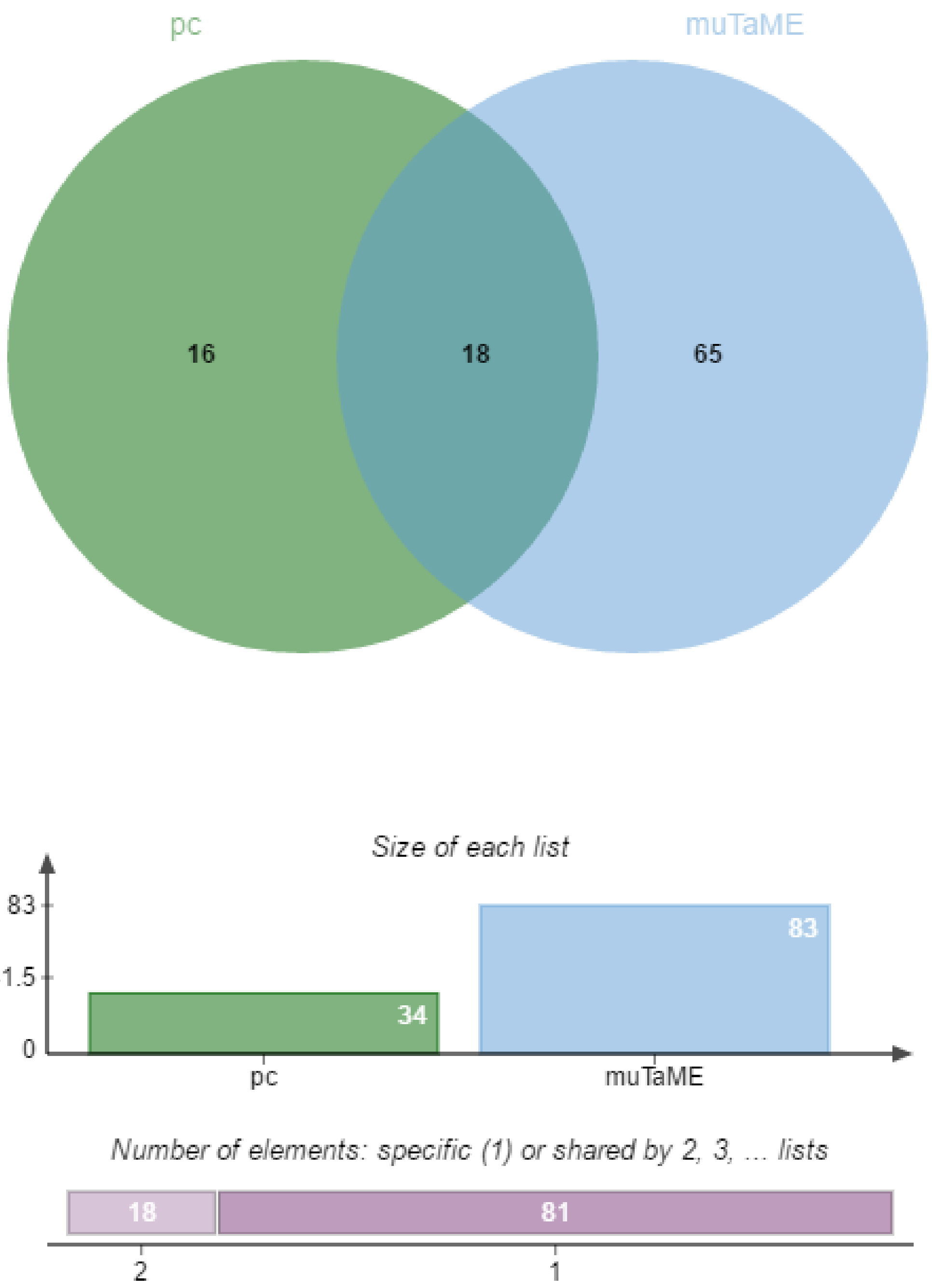
pc
muTaME
16
18
65
Size of each list
83
41.5
0
34
83
pc
muTaME
Number of elements: specific (1) or shared by 2, 3, ... lists
18
81
2
1

K

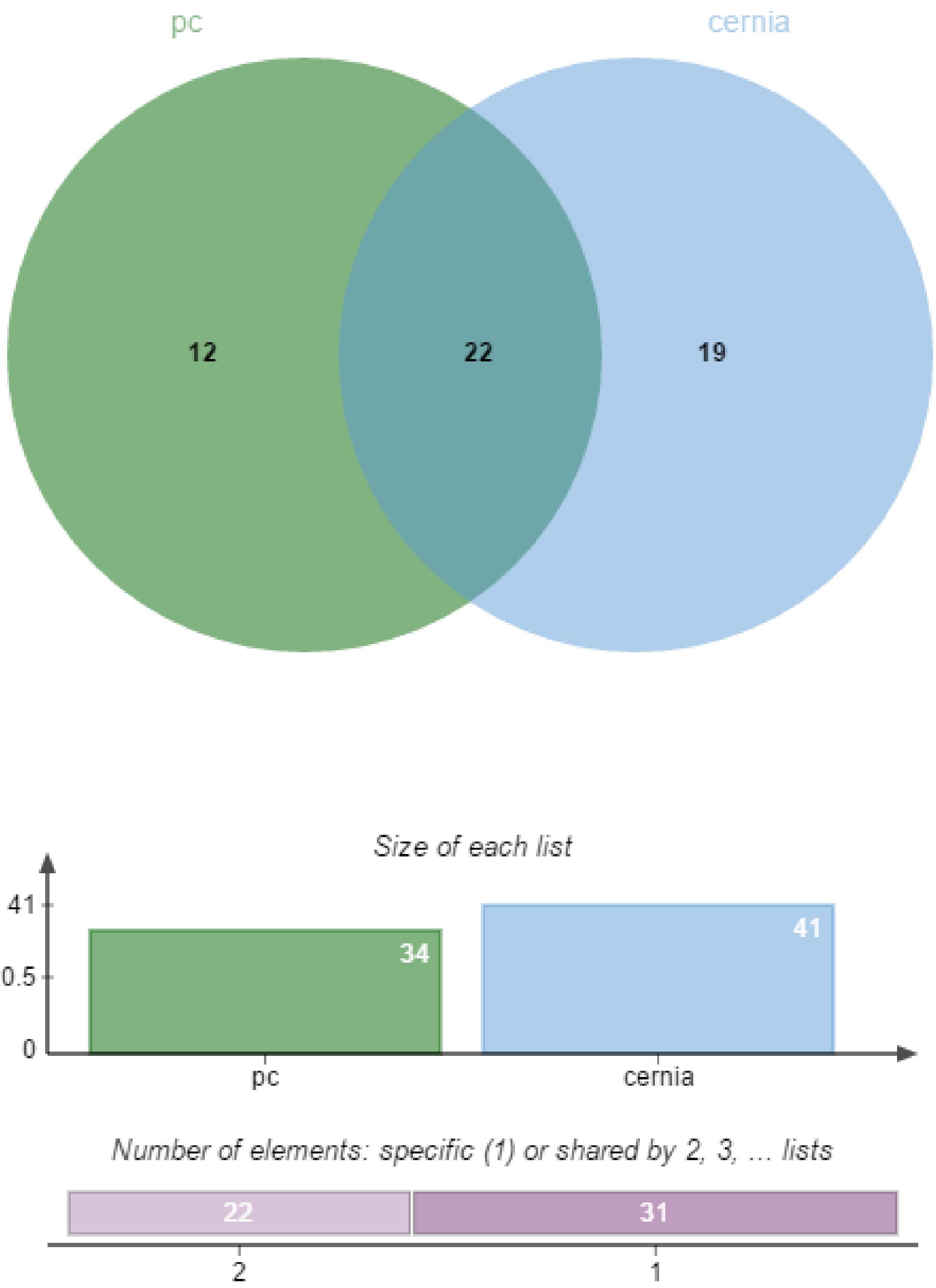
pc
cernia
12
22
19
Size of each list
41
20.5
0
34
41
pc
cernia
Number of elements: specific (1) or shared by 2, 3, ... lists
22
31
2
1

L

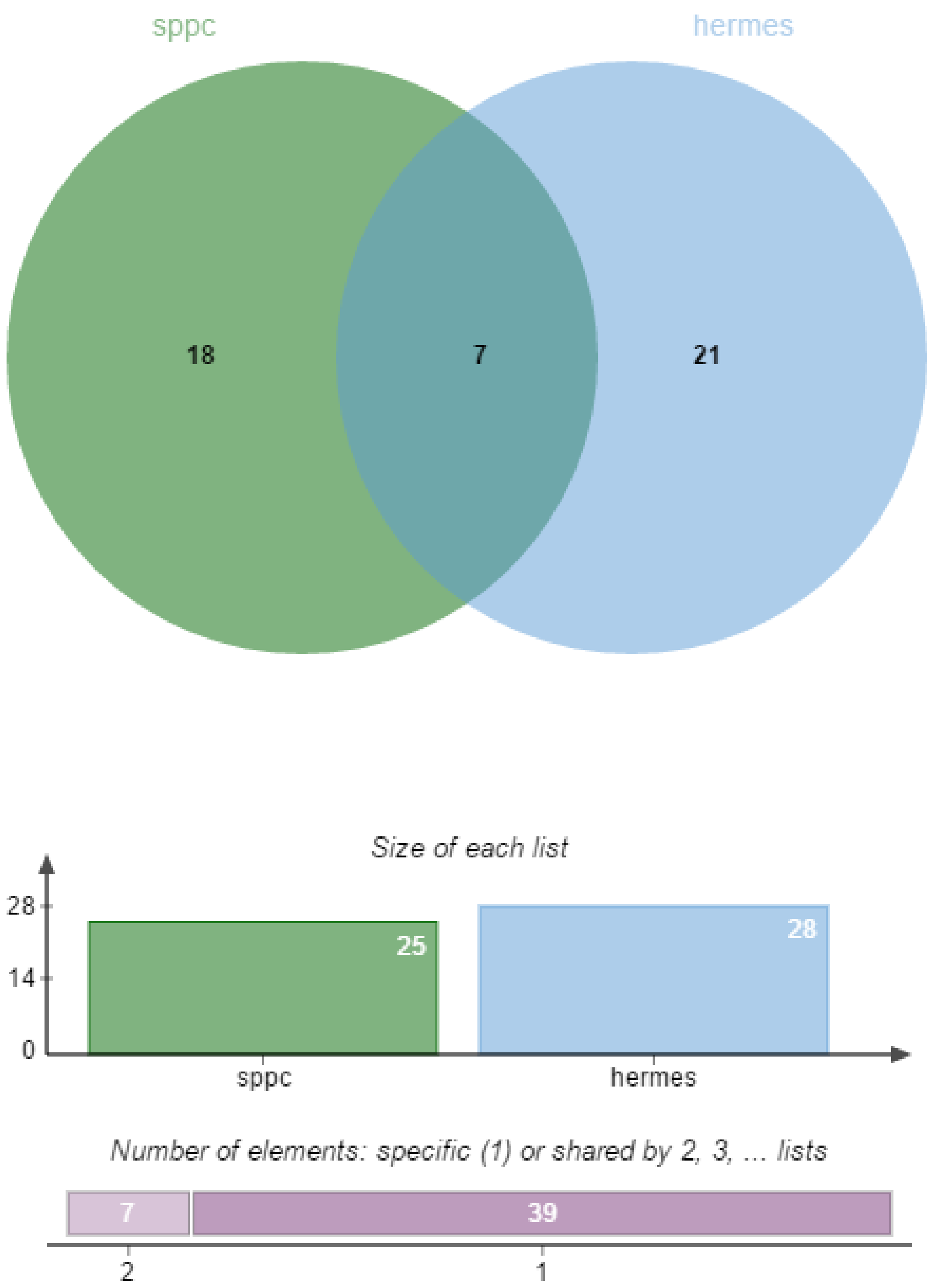

sppc
hermes
18
7
21
Size of each list
28
14
0
25
28
sppc
hermes
Number of elements: specific (1) or shared by 2, 3, ... lists
7
39
2
1

M

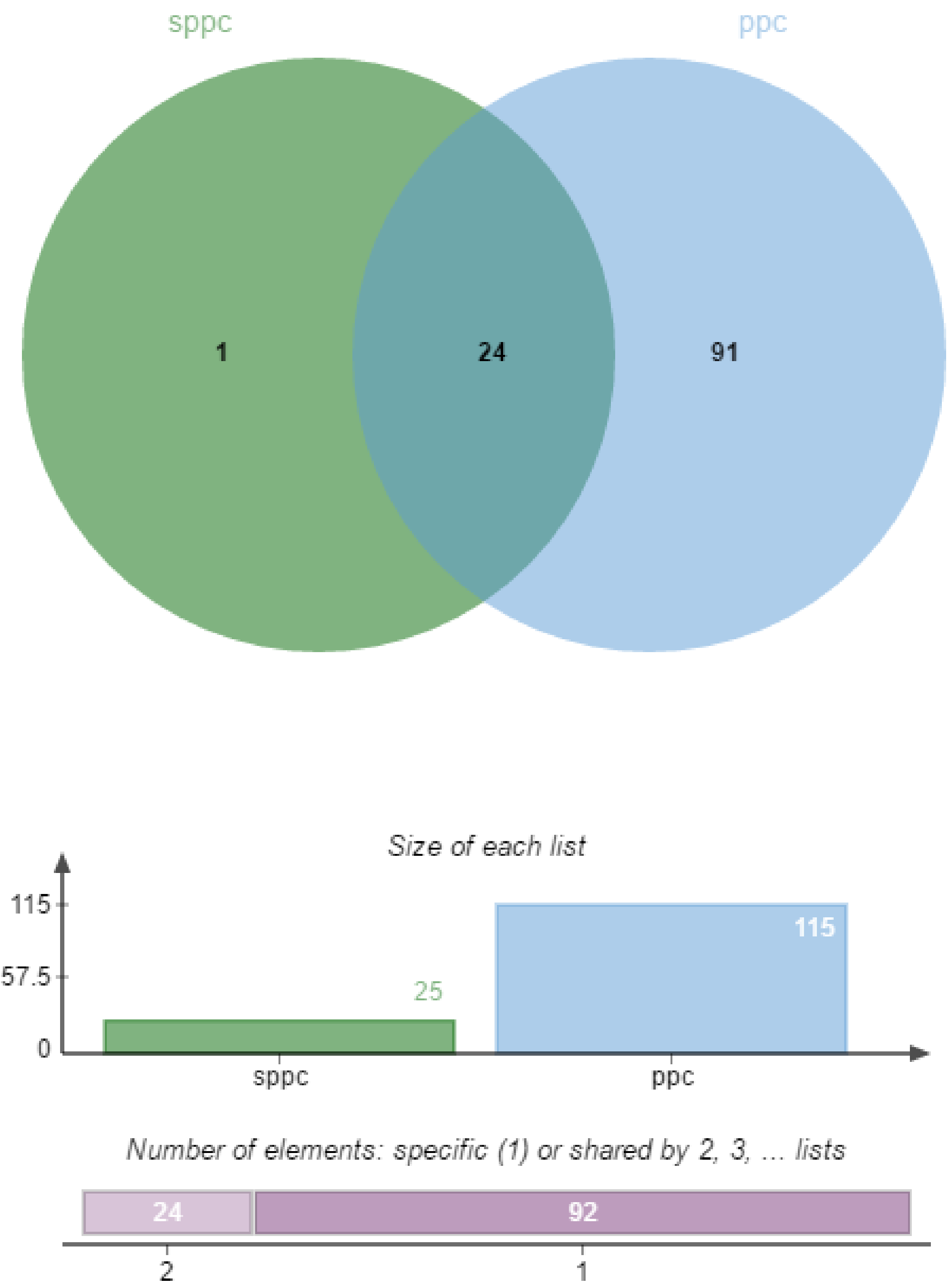
sppc
ppc
1
24
91
Size of each list
115
57.5
0
25
115
sppc
ppc
Number of elements: specific (1) or shared by 2, 3, ... lists
24
92
2
1

N

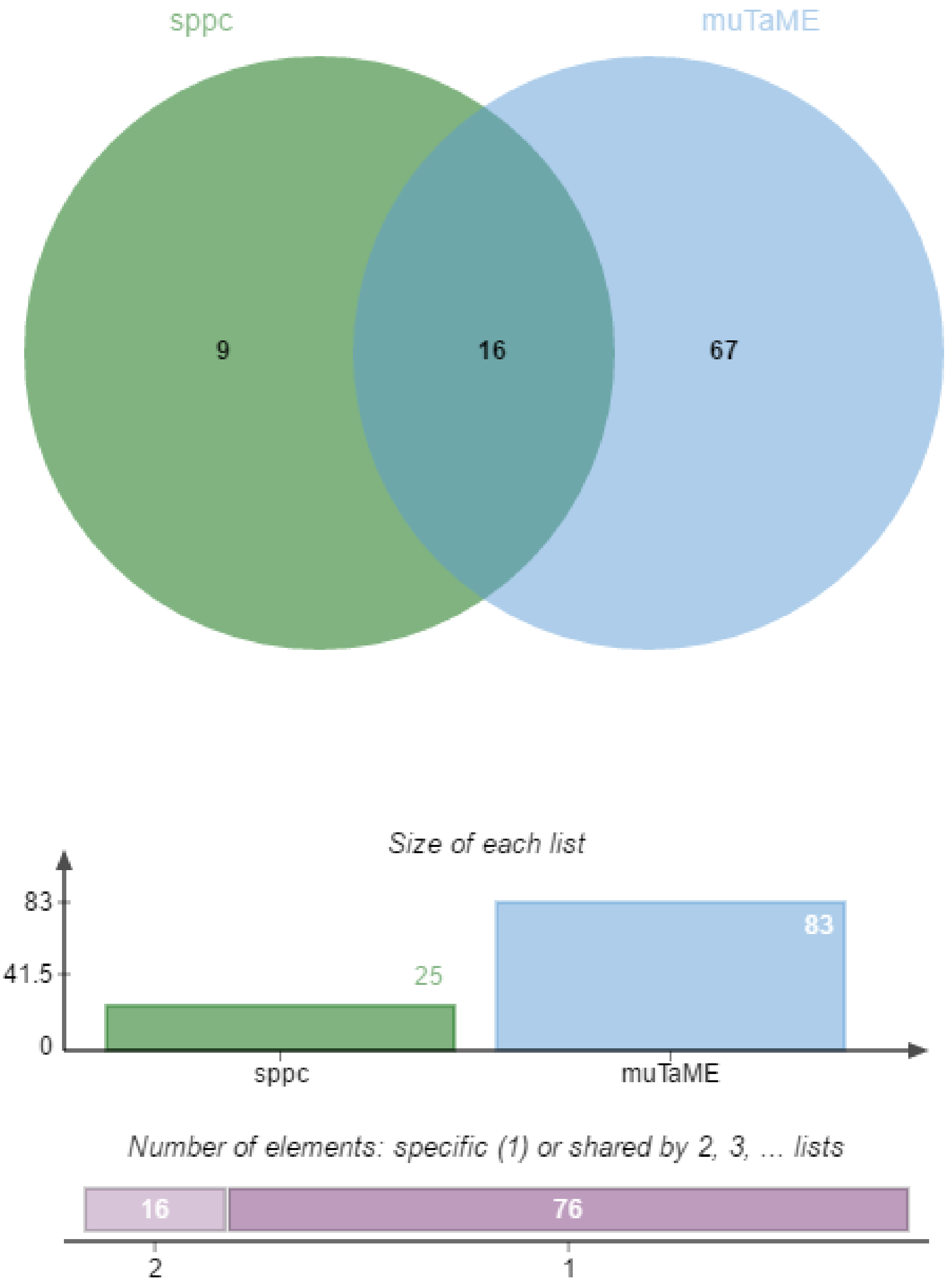
sppc
muTaME
9
16
67
Size of each list
83
41.5
0
25
83
sppc
muTaME
Number of elements: specific (1) or shared by 2, 3, ... lists
16
76
2
1

O

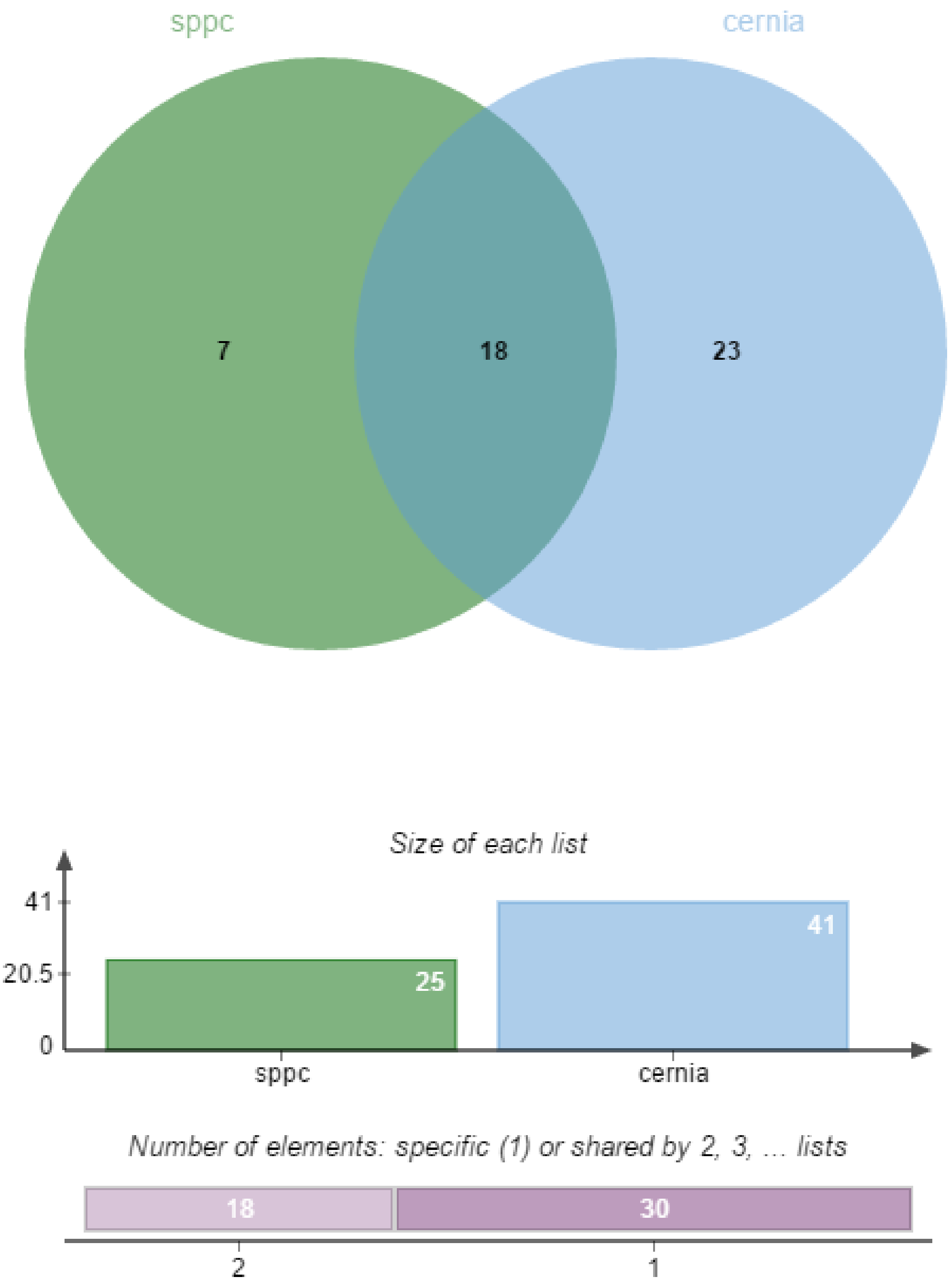
sppc
cernia
7
18
23
Size of each list
41
20.5
0
25
41
sppc
cernia
Number of elements: specific (1) or shared by 2, 3, ... lists
18
30
2
1

P

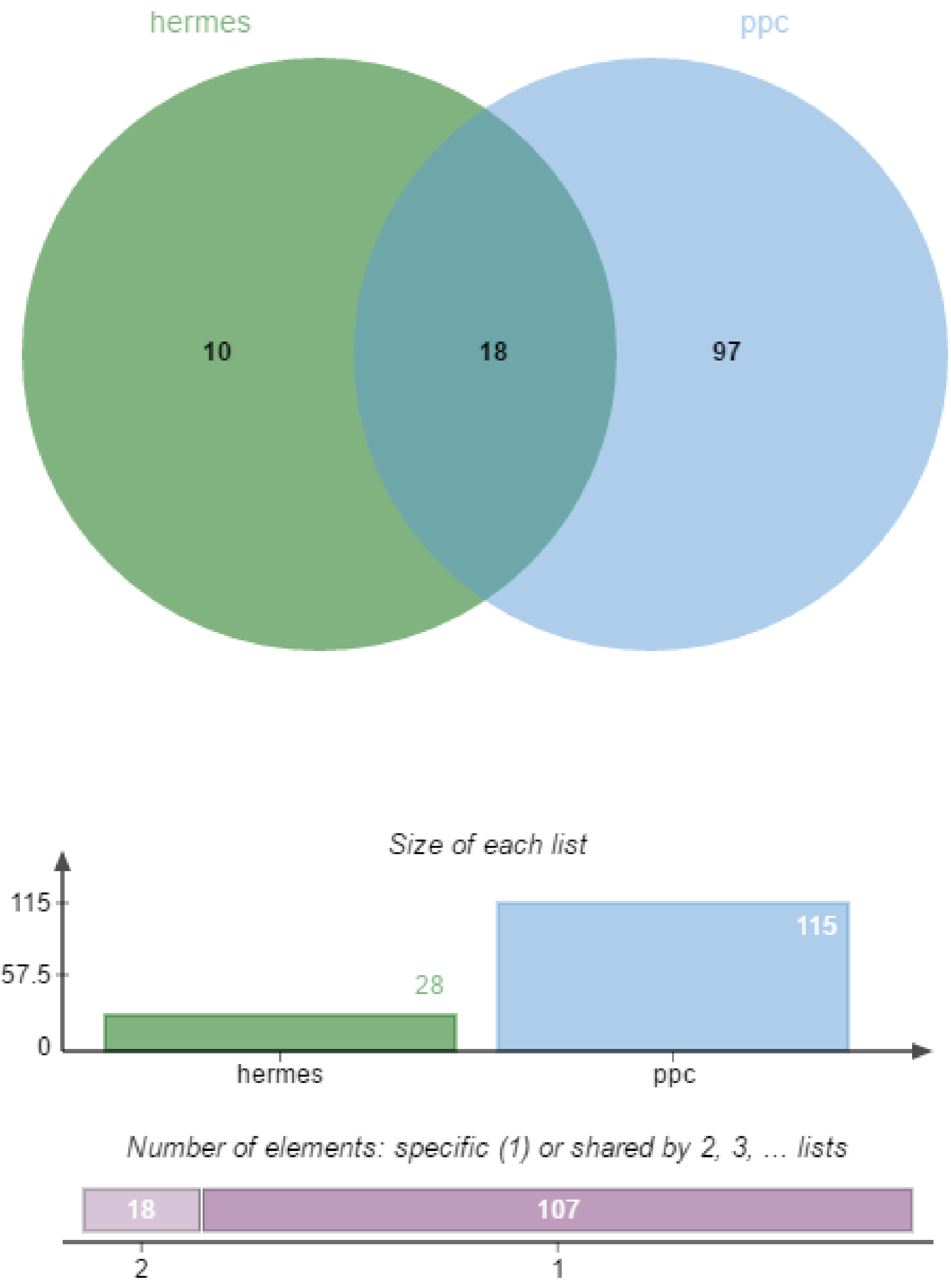
hermes
ppc
10
18
97
Size of each list
115
57.5
0
28
115
hermes
ppc
Number of elements: specific (1) or shared by 2, 3, ... lists
18
107
2
1

**Q**

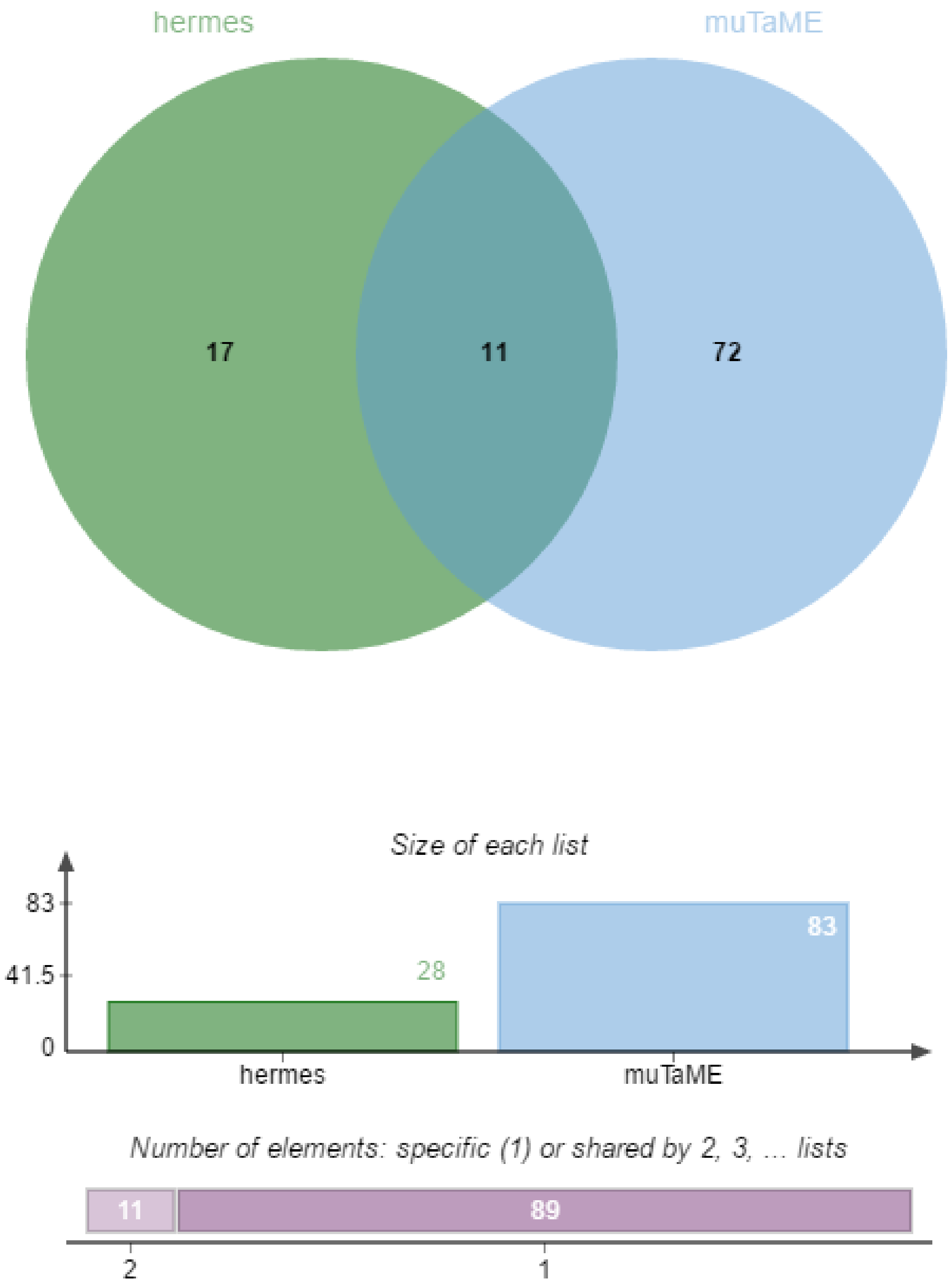
hermes
muTaME
17
11
72
Size of each list
83
41.5
0
28
83
hermes
muTaME
Number of elements: specific (1) or shared by 2, 3, ... lists
11
89
2
1

**R**

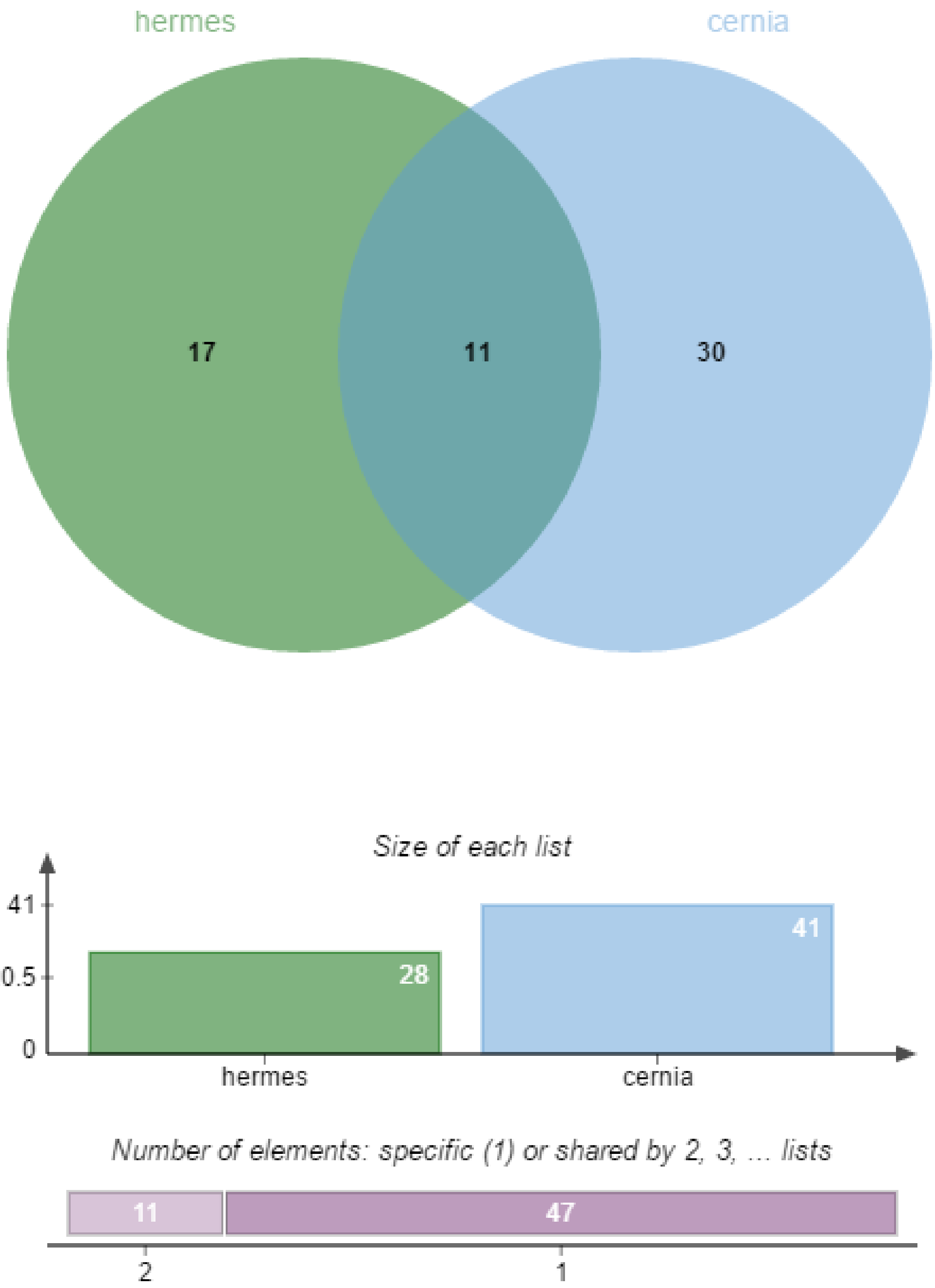
hermes
cernia
17
11
30
Size of each list
41
20.5
0
28
41
hermes
cernia
Number of elements: specific (1) or shared by 2, 3, ... lists
11
47
2
1

S

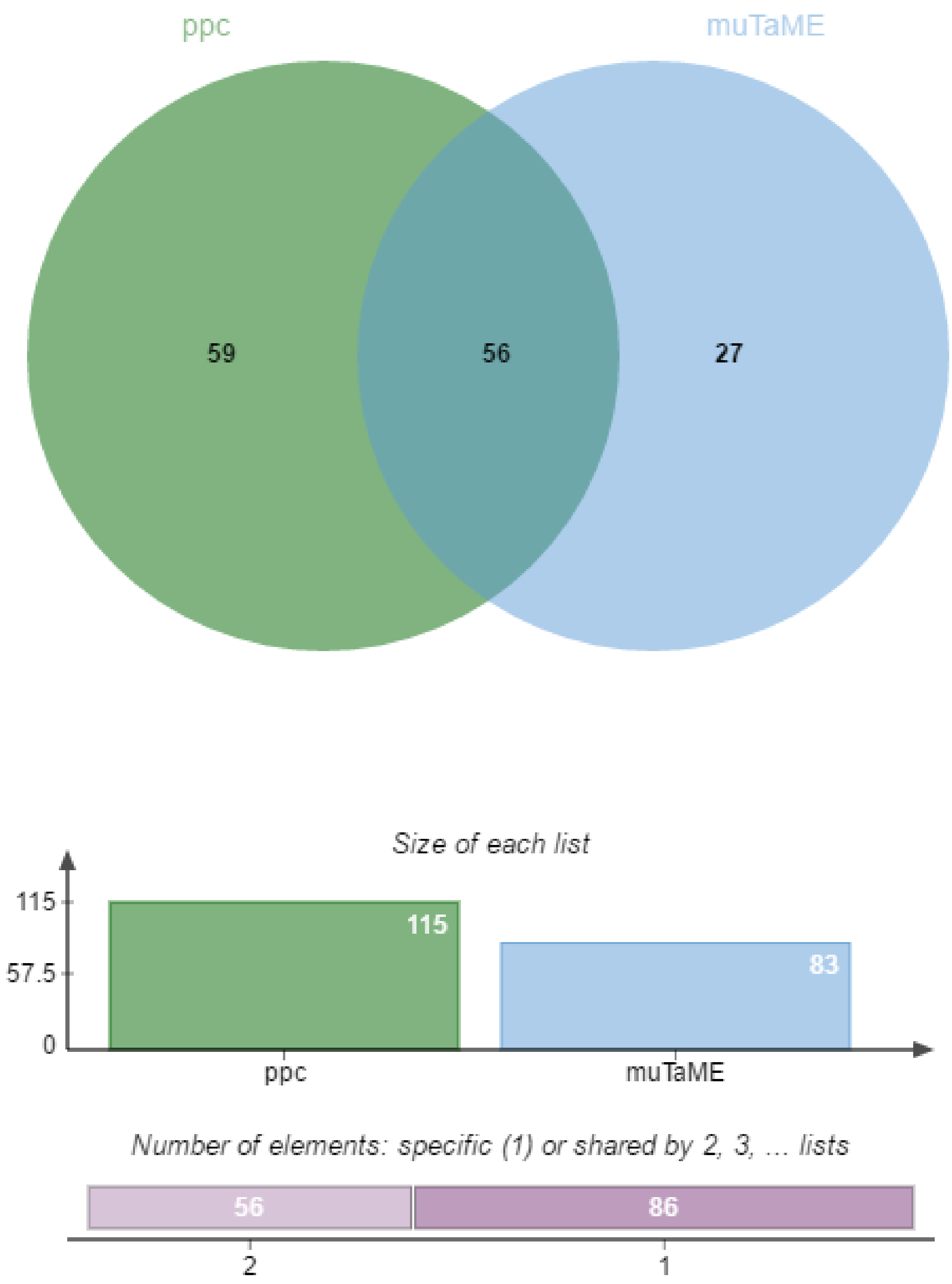

ppc
muTaME
59
56
27
Size of each list
115
57.5
0
115
83
ppc
muTaME
Number of elements: specific (1) or shared by 2, 3, ... lists
56
86
2
1

T

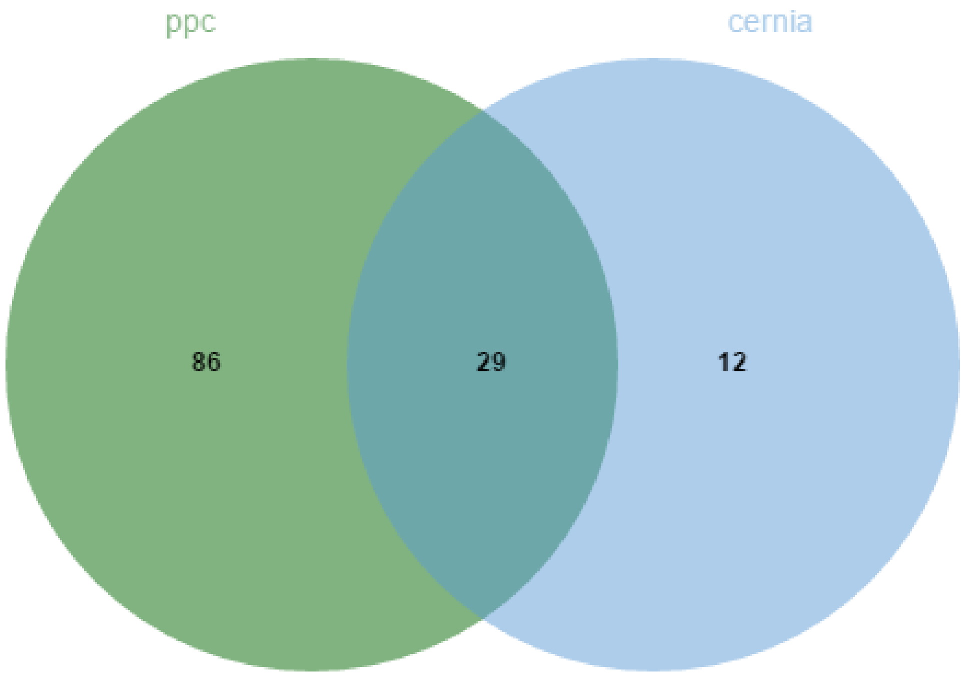
ppc
cernia
86
29
12

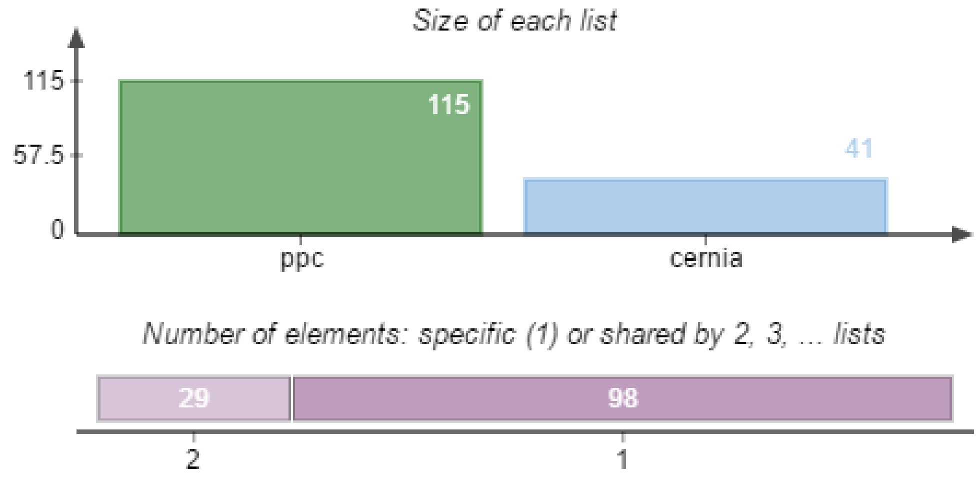
Size of each list
115
57.5
0
115
41
ppc
cernia
Number of elements: specific (1) or shared by 2, 3, ... lists
29
98
2
1

U

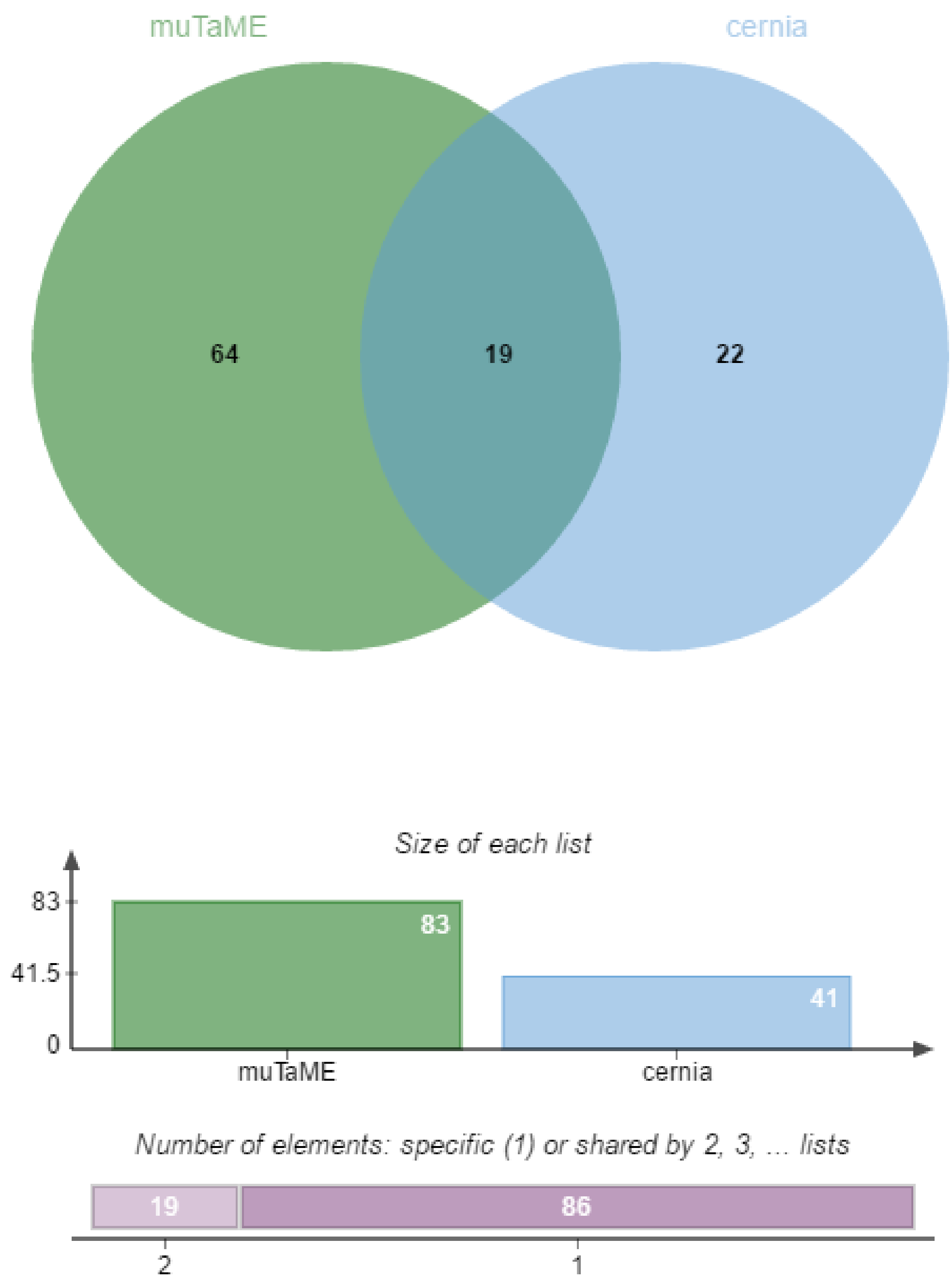


**Figure S1. Pair-wise comparison of overlapping results for 7 individual methods.**

The Venn diagrams are generated by the online tool jvenn (http://bioinfo.genotoul.fr/jvenn).
